# On metallic luster and iridescence in animal coloration

**DOI:** 10.1101/2023.10.12.562066

**Authors:** Klara K. Nordén, Raphael S. Steiner, Anna B. Stephenson, Mary Caswell Stoddard

## Abstract

Some structural colors in nature are frequently described as metallic. For example, hummingbird plumage, jewel beetles and *Morpho* butterflies have this attribute. While much attention has been paid to describing the often-shifting hues of these structural colors, there has been little interest in explaining why they appear metallic. In this paper, we argue that the metallic luster (the metallic appearance or sheen) of some structural colors arises in part from a combination of two factors: a colored specular reflection and a very low diffuse reflection. Reflections with these characteristics are found in metals and are distinct from other material reflections in nature. We propose that metallic luster can be classified based on these two reflectance properties (colored specular reflection and low diffuse reflection). We also suggest that some of the ambiguity surrounding the term “iridescent structural color” can be traced to the frequent confounding of metallic luster with a common definition of iridescence: a shift of peak spectral wavelength (often referred to as hue) with viewing angle. We show using optical models and cross-polarization imaging of bird plumage that two types of structural colors that are often classified as “iridescent” and “non-iridescent” both display iridescence—but only one type has metallic luster. By considering metallic luster and iridescence separately, we simultaneously clarify terminology in structural colors and open up many new lines of inquiry regarding the perception of metallic luster in animals.

## 1. Introduction: changeable colors

In many cultures, “iridescence” has been used to symbolize a mysterious or unpredictable power that is difficult to grasp (Sutton & Snow, 2015). In a curious parallel, the term “iridescence” in natural sciences is difficult to define and is used to describe a range of visual effects. “Iridescence” has been used to describe rainbow-like colors in beetles and flowers (Seago et al., 2009), the pearlescent colors of nacre (Ozaki et al., 2021), and the metallic colors found in, for example, bird plumage and insect cuticles (Doucet & Meadows, 2009, Seago et al., 2009). In The Oxford English Dictionary, the entry for “iridescent” reads: “displaying colours like those of the rainbow, or those reflected from soap bubbles and the like; glittering or flashing with colours which change according to the position from which they are viewed” (Simpson et al., 1989). This definition thus involves rainbow-like colors, colors that change with viewing angle, and glittering or flashing of colors (similar to metallic luster). However, all three of these effects do not necessarily occur together (for example, the rainbow is not glittering or flashing but displays rainbow colors which change with viewing angle). Therefore, a more precise definition of “iridescence” is needed in the sciences.

This point has been emphasized in previous reviews of “iridescence” in nature, which address the problem by restricting the definition of iridescence to “a change in hue with viewing or illumination angle” (Stuart-Fox et al., 2020, Ospina-Rozo et al., 2022). Stuart-Fox et al. (2020) and Ospina-Rozo et al. (2022) use the word hue as opposed to the broader term color (which includes hue, saturation and brightness), since almost all objects will change in brightness and saturation with viewing or illumination angle due to the effects of specular reflection. To get an intuition for this, imagine turning a shiny, red apple. As you do so, specular highlights will move across the surface of the apple. Thus, the brightness and saturation of the apple change with viewing and illumination angle. To exclude such effects, recent reviews of “iridescence” in nature have therefore argued that the term “iridescence” should be restricted to changes in hue (Stuart-Fox et al., 2020, Ospina-Rozo et al., 2022), while gloss is an already well-established term for angle-dependent changes in brightness and saturation (Chadwick & Kentridge, 2015). Following these reviews (Stuart-Fox et al., 2020, Ospina-Rozo et al., 2022), we use the term iridescence to mean a shift in peak spectral wavelength with a change in observation or viewing geometry, while “iridescence” (in quotation marks) refers to the concept in a broader sense (see §2, *Definitions*). With the stricter definition of iridescence, it might appear that the confusion surrounding the term “iridescence” has been resolved. However, we argue that “iridescence” is still problematic since many authors still use it to describe other features of “iridescence” that have not been properly characterized and defined. In this paper, we characterize one such usage in the bird coloration literature and give it a name: *metallic luster*.

Metallic luster is a term originating from geology, where it is defined as “having the sheen characteristic of metal” (Allaby, 2013). Minerals with metallic luster include pure metals such as gold, copper and silver, as well as metal-containing compounds such as pyrite. Many minerals combine iridescence with metallic luster, such as bornite (often called “peacock ore”), yet it is clear that these two aspects of minerals are distinct (Figure 1). It is possible to exhibit iridescence without metallic luster (such as in the rainbow, Figure 1A) and metallic luster without iridescence (such as in gold, Figure 1E). The key argument of our paper is that this dichotomy has remained unappreciated in the animal coloration literature. “Iridescence” is often used to describe objects that exhibit metallic luster or metallic luster in combination with iridescence (e.g. by some of us previously in Nordén et al., 2021). However, “iridescence” is rarely used to describe iridescent objects that lack metallic luster. This has two unfortunate consequences. First, the iridescence of objects that lack metallic luster has been explored little and is generally disregarded. Second, for objects that exhibit both iridescence and metallic luster, only the iridescent qualities are emphasized, with little attention to metallic luster. To rectify these problems, we will show that many structural colors in nature that are referred to as “non-iridescent” are in fact iridescent, just like structural colors often referred to as “iridescent”. We will also introduce a method for approximating a material’s metallic luster, offering researchers a way of describing the distinct contribution of metallic luster to structural colors. We hope that this will inspire new questions in the study of these remarkable colors.

**Figure 1.**
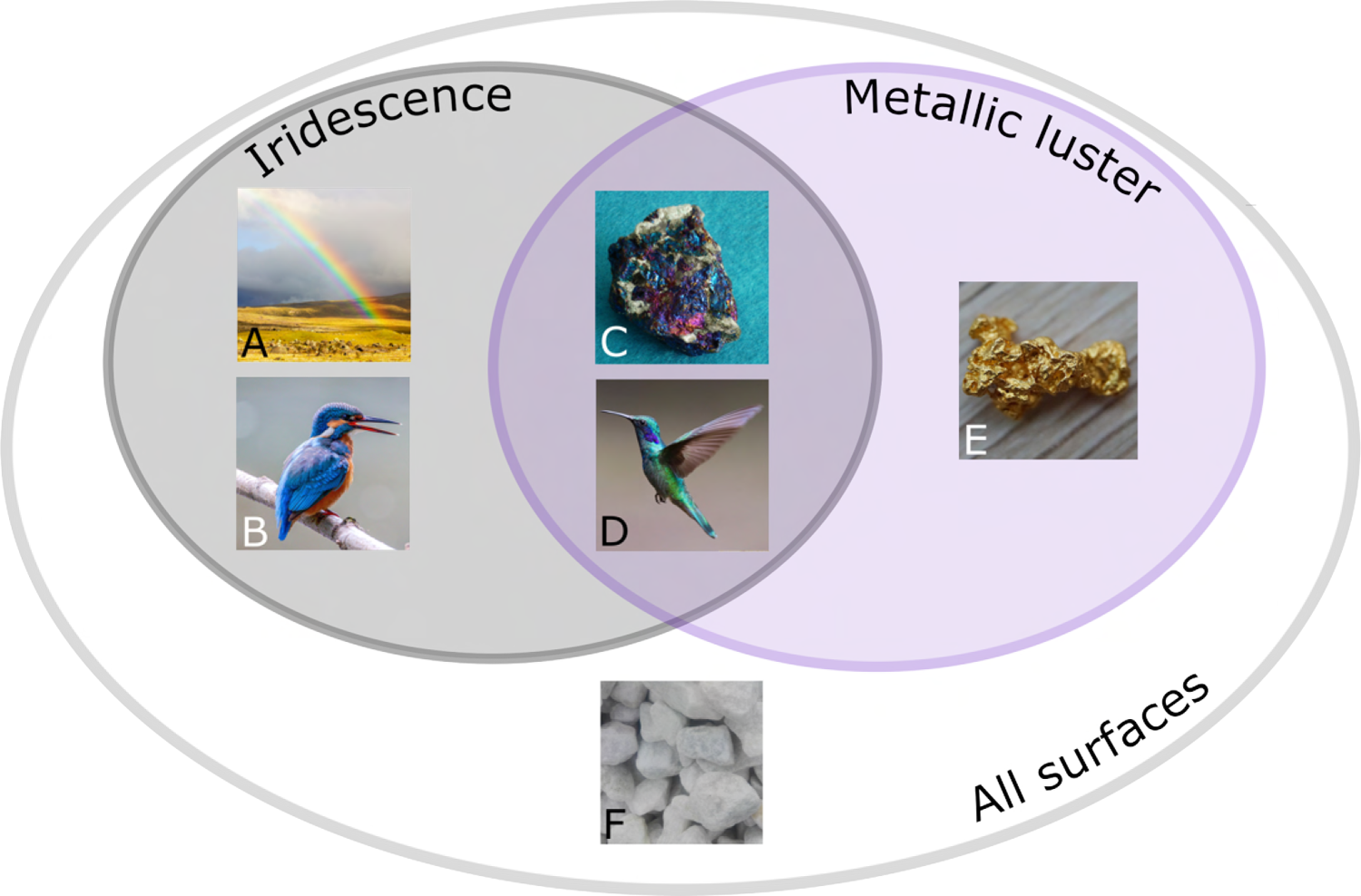
Iridescence and metallic luster are two distinct properties that are often confused. Even though some objects are both iridescent and exhibit metallic luster, such as bornite (C) and the plumage of a hummingbird (D), these two properties need not be linked. Objects can be iridescent and lack metallic luster, as is the case for the rainbow (A) and the blue plumage of a Common kingfisher (*Alcedo atthis*, B), or exhibit metallic luster but lack iridescence, as is the case for gold (E). Many objects, of course, lack both features—for example calcium carbonate (F). Images sourced from pixabay.com under a Pixabay license.

The paper is structured as follows: In §2, *Definitions*, we define various terms we will use throughout the paper.

In §3, *Birds of an iridescent feather*, we use examples from the bird coloration literature to illustrate that two types of structurally colored plumage, commonly called “iridescent” and “non-iridescent”, do not in fact differ in iridescence when measured over changing angles between illumination and observation. We support our argument with a theoretical analysis of peak spectral wavelength shifts for both “iridescent” and “non-iridescent” plumage, as well as with data from the literature of measured peak spectral wavelength shifts in both types of structurally colored plumage.

In §4, *All that glitters is not gold*, we outline what a metallic reflection is and give a verbal argument based on the known optics of feather nanostructures that can explain why they—despite not containing any metals—appear metallic. We argue that it is in fact this metal-like reflection (metallic luster) that clearly sets “iridescent” structural colors apart from other structural and pigmentary color mechanisms, and we propose a classification based on two reflection properties that are characteristic of objects with metallic luster.

In §5, *Measuring metallic luster*, we develop a simple quantitative measure of metallic luster based on these reflectance properties using cross-polarization photography. We demonstrate this method using a sample of plumage patches with different color mechanisms.

Lastly, in the *Discussion*, §6, we discuss the merits of incorporating metallic luster into discussions of animal structural color. In particular, we argue that doing so is more than an exercise in building precise definitions: it in fact opens the door to intriguing research questions and new directions.

## 2. Definitions

Due to the aforementioned ambiguity surrounding the term “iridescence”, we will give precise definitions of *iridescence*, *metallic luster*, and related terms, of which we shall henceforth make use. When relevant, we have added remarks to give additional background and motivation for our usage of the term.

### Hue

The tint of a color, which is dependent on the spectral distribution of the reflectance, the illumination and the viewer’s visual system.

#### Remarks

This term is interchangeably used in the animal coloration literature to describe a perceptual quality (as in our definition), or a proxy for this, usually the peak spectral wavelength (Montgomerie, 2006). The measures are related—but differ fundamentally since perceptual hue depends on the viewer’s visual system, while peak spectral wavelength is an objective measure from the physical reflectance spectrum.

### Peak spectral wavelength

The wavelength of maximum reflectance in a reflectance spectrum.

#### Remarks

There are many variations of this measure suited to different types of spectra. For example, spectral location (Ospina-Rozo et al., 2022) may better capture variations of broad spectral peaks. However, all such measures aim to specify the position of the spectral peak and are thus conceptually the same. Since the spectra we are analyzing in this paper generally have well-defined peaks, we used peak spectral wavelength.

### Iridescence

A shift in peak spectral wavelength with change in observation or viewing geometry.

#### Remarks

More commonly, this is defined as a shift in hue with viewing or observation angle, but with the dual meaning of hue implied (perceptual or objective) it can thus be interpreted as either a perceptual or an objective property. Since iridescence is almost always measured as a shift in peak spectral wavelength (or some version thereof) and this measure does not require choosing a specific visual system, we use peak spectral wavelength to develop our argument in this paper.

#### “Iridescence”

Indicates general uses of the term, including rainbow colors, pearlescence (appearance similar to a pearl), glossiness and metallic luster, and/or where the precise meaning was not defined in the source referenced.

### Metallic luster

Metallic sheen of an object, the reflection of which can be characterized as having

1. a colored specular reflectance; and
2. very low diffuse reflectance.

#### Remarks

Metallic luster is a perceptual concept, which is difficult to measure directly. However, it can be measured by a proxy. In this paper, we use aspects of the specular and diffuse reflection, as defined above.

### Specular reflectance

A reflection seen at the mirror angle of light incidence with respect to the tangent of the surface (i.e. angles of light incidence and reflection are equal).

#### Remarks

On rough surfaces, the specular reflection will appear blurry due to the influence of the varied surface geometry (Figure 5B). On smooth surfaces, the specular reflection will appear sharp (**Gloss**, Figure 5A). Note that we have adapted here definitions of specular reflectance and gloss that are typically used in reflection models for computer graphics—see Ginneken et al. (1998).

### Diffuse reflectance

A reflection that is seen at equal intensity from all viewing angles.

#### Remarks

Diffuse reflectance arises from sub-surface scattering in a material. This causes the light to be scattered multiple times, and it will therefore exit the material at a random angle. Note that a similar effect can be produced by the specular reflection from a rough surface (Figure 5B). By some authors, surface reflection from rough surfaces is therefore described as “diffuse”. Here, with diffuse reflection we mean sub-surface scattering only, following definitions of diffuse and specular reflectance that are typically used in reflection models for computer graphics—see Ginneken et al. (1998).

### Single-scattered light

Light that has only been scattered a single time.

#### Remarks

Specular reflections originate from single-scattered light. Note that while a specular reflection is defined by its direction (reflected at the mirror angle), the term single-scattered is only defined by the type of scatter, not its direction.

### Multiple-scattered light

Light that has been scattered multiple times.

#### Remarks

Diffuse reflections of objects arise from multiple-scattered light. Note that while a diffuse reflection is defined by the uniform distribution of the reflectance direction, the term multiple-scattered light is only defined by the type of scatter, not its direction.

### Gloss

The specular reflection of a smooth object.

#### Remarks

For most objects, the gloss will have the same spectral distribution as the illumination (typically white). The exceptions are metals and some structurally colored objects, which can reflect colored gloss.

### Structural color

Colors arising from the interaction of light with a structure (i.e. a material with spatial inhomonogeneity). Optical processes such as reflection, refraction, scattering, interference and diffraction are the mechanisms behind structural color production.

#### Remarks

Structural colors can be contrasted with pigmentary colors, where the color arises from the interaction of light with an absorbing pigment. Note however, that many colors in nature involve both mechanisms, but to different degrees.

### Barbule structural color

Type of structural colors in feather barbules, consisting of ordered arrays of melanin granules (melanosomes, Figure 2A). In an optical sense, these structures can be approximated well by a multilayer structure. In the context of bird feather colors, often used interchangeably in the literature with *“iridescent” structural colors*. To our knowledge, barbule structural colors typically exhibit both iridescence and metallic luster (but see §Appendix A).

**Figure 2.**
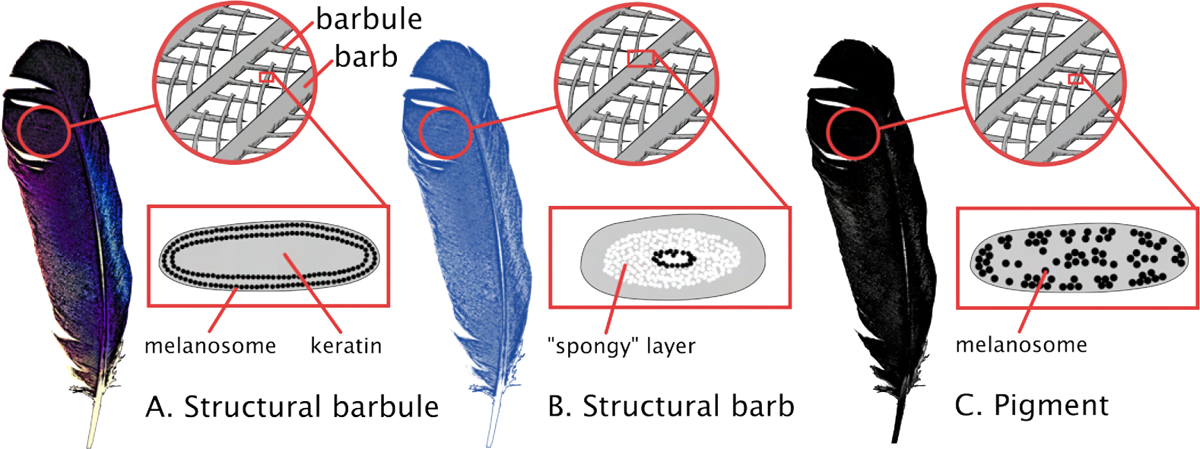
The color mechanisms of feathers can be divided into three main categories: structural barbule (A), structural barb (B) and pigmentary (C). Cross-sections of barbs/barbules shown in boxes. Structural barbules (A) give rise to shimmering and metallic colors and are produced from ordered arrays of melanin-containing organelles (melanosomes) in the barbules. Structural barbs (B) give rise to diffuse blue and green colors and are produced from a sponge-like structure in the barbs made of keratin and air underlain by melanin. Pigmentary colors (C) give rise to a range of different hues depending on the absorbance of the pigment. The pigment is usually distributed in both barb and barbules. Here, a black melanin-colored feather is pictured with disordered melanosomes in the barbules (compare to (A)).

### Barb structural colors

Type of structural colors in feather barbs, consisting of air and keratin (spongy keratin, Figure 2B). In an optical sense, these structures can be approximated well by a photonic glass. In the context of bird feather colors, often used interchangeably with *“non-iridescent” structural colors*. To our knowledge, barb structural color typically exhibit iridescence but never metallic luster.

## 3. Birds of an iridescent feather

To illustrate our argument that the term “iridescent” is often confounded with metallic luster, we will use bird plumage coloration, since there is a long history of classifying structural coloration of feathers as “iridescent” and “non-iridescent” (Gadow, 1882, Auber, 1957, Michelson, 1911, Haecker, 1890). In birds, one type of structural color is located in the feather barbules, gives a metallic shine and tends to shimmer when viewing angle is varied (Figure 2A). This type of coloration is found in, for example, the European starling (*Sturnus vulgaris*), Indian peafowl (*Pavo cristatus*) and many hummingbirds (family Trochilidae). A second type of structural color is located in the feather barbs and appears diffuse—not unlike pigmentary colors (Figure 2B). This type of coloration gives rise to, for example, the blue of the European kingfisher (*Alcedo atthis*), Blue jay (*Cyanocitta cristata*) and Indigo bunting (*Passerina cyanea*). Thus, there is clearly a difference in the visual appearance of these two types of colors (at least to humans). Though they are commonly called “iridescent” and “non-iridescent”, what we aim to show is that iridescence is in fact not the main visual property in which they differ. We stated in the previous section that previous authors have suggested that angle-dependent change in reflectance can be divided into two terms: gloss (or a change in intensity/spectral purity with viewing angle) and iridescence (a change in peak spectral wavelength with viewing angle) (Stuart-Fox et al., 2020, Seago et al., 2009, Ospina-Rozo et al., 2022). However, neither of these properties, gloss and iridescence, separates structural barbule coloration (“iridescent” structural colors) from barb coloration (“non-iridescent” structural colors). We unpack this in the sections below.

### 3.1. Gloss

The first aspect of angle-dependent change is gloss, which involves a change in light intensity with angle. Gloss appears as a white highlight on most objects, but can be colored for some structurally colored objects and metals. All structural barbule coloration is glossy—but not all glossy plumage is produced by structural barbule coloration. Gloss is frequently found in pigmentary plumage (Iskandar et al., 2016) as well as plumage with structural barb coloration (Figure 3). One might imagine that glossiness (measured as a change in reflected intensity with angle) is greater in structural barbule colors than in pigmented or structural barb colors, but this is not true. While some structural barbule coloration can achieve a reflectance ranging from almost zero to 100% when viewing/illumination angle is varied, the same is also true for glossy dark pigmentary plumage (a dark surface will reflect almost no light if illuminated perpendicularly, but nearly 100% when illuminated almost parallel to the surface, as described by the Fresnel equations). Thus, neither the presence nor magnitude of gloss uniquely captures structural barbule coloration—despite gloss being a feature of this type of coloration.

**Figure 3.**
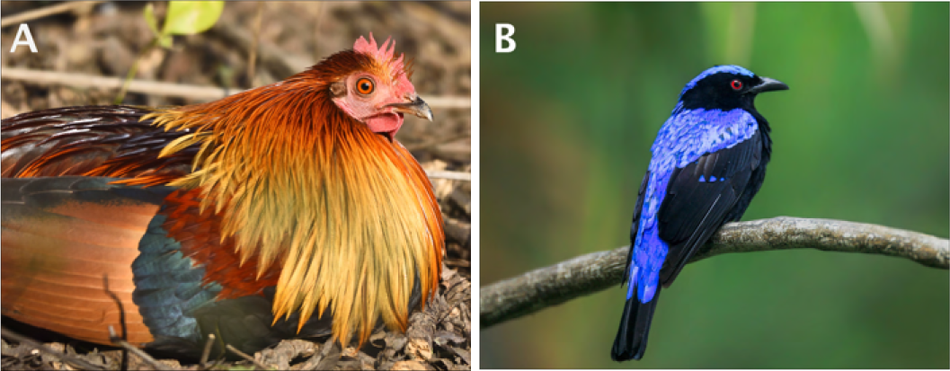
Gloss is a common property of many types of plumage, including pigmentary and structural barb colors. A) Glossy pigmentary feathers on the back of the Junglefowl (*Gallus gallus*). B) Glossy structural barb coloration in the Asian fairy blue-bird (*Irena puella*). Image credits: A) all rights reserved by copyright holder Praveen Pandian, reproduced here with permission; B) all rights reserved by copyright holder Akhanhk, reproduced here with permission.

### 3.2. Iridescence

Now, let us turn to the second aspect of angle-dependent change— iridescence. It turns out that “iridescent” and “non-iridescent” structural colors are similar in presence and degree of iridescence. This statement is quite surprising, and we will explore it in more depth using a theoretical approach which we will validate with measurements of plumage taken from published studies.

Taking a theoretical approach to demonstrate that structural barbule and structural barb coloration do not necessarily differ in presence or magnitude of iridescence, we modeled structural barbules as multilayer structures and structural barbs as photonic glasses, since this approximates the optics of the real structures well (Kinoshita et al., 2008, Stavenga et al., 2017, Prum, 2006). There are well-known relationships for how multilayers and photonic glasses scatter light with varying angle of incidence, which we adapted here to model iridescence (Kinoshita et al., 2008, Magkiriadou et al., 2014). Since multiple measurement geometries are possible to quantify iridescence, we chose the geometry that captures the maximum peak spectral wavelength shift of a multilayer (“constant angle bisector” *sensu* Gruson et al., 2019), where the reflectance is measured at the mirror angle of light incidence (i.e. a specular configuration). We searched a parameter space that is relevant to bird plumage, using ranges derived from previously published studies of feather nanostructures. Our model estimates of structural barbules and barbs are simplifications, since structural barbules are not perfect multilayers and structural barbs can often display higher nanostructural ordering than a photonic glass. Note, however, that our simplifying assumptions bias our estimates of iridescence towards more strongly angle-dependent barbule structural colors and less angle-dependent structural barb colors—i.e. it is a conservative estimate because we might expect barbule structural colors to be more angle-dependent than structural barb colors.

#### 3.2.1. Optical modeling and plumage data

The iridescence of multilayer structures (structural barbules) is estimated using the relationship:

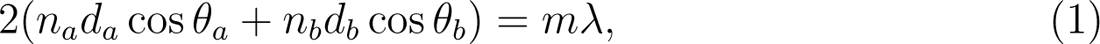

where *n_a_* and *n_b_* are the refractive indices of the two materials in the multilayer, *d_a_* and *d_b_* are the thicknesses of the layers, and cos *θ_a_* and cos *θ_b_* are the angles of refraction in the materials (see Kinoshita et al., 2008 for a review of multilayer optics). We used refractive indices of 1.75 and 1.58 for melanin and keratin, respectively (Stavenga et al., 2015). The thicknesses of the layers were taken from a database of all known multilayer structures in feathers (Nordén et al., 2021), which was filtered to include only structures with melanin and keratin.

The iridescence of photonic glasses (structural barbs) is approximated by the relationship

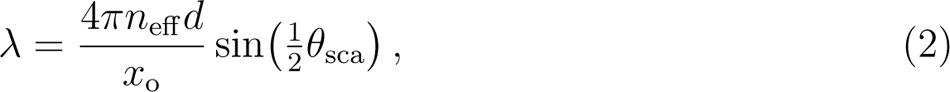

where *λ* is the peak spectral wavelength, *n*_eff_ is the effective refractive index, *d* is the diameter of particles, and *θ*_sca_ is the scattering angle (Magkiriadou et al., 2014). Further, *x*_o_ is the magnitude of the dimensionless wavevector at the peak in the structure factor, defined as *x*_o_ = (2*πd*)*/*(*d*_avg_), where *d*_avg_ is the average center-to-center spacing between coordination shells of particles (Magkiriadou et al., 2014). In the disordered porous packing in bird feathers, coordination shells are the arrangements of pores surrounding an arbitrary central pore. The first coordination shell refers to the pores directly in contact with the central pore. The second coordination shell refers to pores in contact with those in the first coordination shell, and so on. We estimate *d*_avg_ to be 0.9d, slightly larger than the distance between FCC (111) planes, which is the plane that scatters most strongly in a face-centered cubic lattice. Since the FCC packing fraction is 0.74 and these disordered structures have a lower packing fraction, we expect the distance between coordination shells to be larger than the distance between FCC (111) planes. We used the Bruggeman approximation to calculate the effective refractive index of structural barbs, using a range of air volume fraction from 50-66%. This range was based on a survey of structural barb structures in 320 species by Saranathan et al. (2012). We varied the diameter of particles over the range 160-250nm, based on values in the literature (Urquia et al., 2020, Prum et al., 2009). We note that our approximation of *λ* (the peak spectral wavelength) is agnostic to the precise packing of the structures. That is, our calculation assumes that the structures are isotropic and have long-range disorder; we do not require a specific structure factor, such as the Percus–Yevick structure factor normally used to describe glasses. However, we still refer to the structures as photonic glasses because Equation (2) captures the peak spectral wavelengths of glasses (as well as other similar structures), and these porous keratin structures in birds are often referred to as photonic glasses in the literature. A summary of all model parameters can be found in Table 1.

**Table 1.** Parameters used for optical modeling.

| Multilayer |  | Photonic glass |  |
| --- | --- | --- | --- |
| Parameter | Value(s) | Parameter | Value(s) |
| $n_a$ | 1.58 | $n_{\text{matrix}}$ | 1.58 |
| $n_b$ | 1.75 | $n_{\text{pores}}$ | 1 |
| $d_a$ | 10-165 nm | $d$ | 160-250 nm |
| $d_b$ | 45-202 nm | $d_{\text{avg}}$ | $0.9d$ |
| | | $vf_{\text{pores}}$ | 0.5-0.66 |

To validate our theoretical models of the iridescence generated by multilayers and photonic glasses, we obtained peak spectral wavelength information from published studies on structural coloration in plumage. We only included studies that used the same geometry for measurement as in our theoretical modeling, i.e. specular configuration. Authors of nine different studies generously shared their plumage data with us, resulting in a data set including 12 species (6 with structural barbule coloration and 5 with structural barb coloration).

#### 3.2.2. Results

The results of our theoretical modeling show that structural barbs (modeled as photonic glass) are capable of producing greater iridescence than structural barbules (modeled as multilayer)—though both types of structures generally overlap (Figure 4A). This result is supported by the plumage measurements we collected from the literature, which show that structural barb and structural barbule colors are broadly overlapping in iridescence (Figure 4B). Moreover, the wavelength shifts observed in our modeling simulations are of a similar magnitude to those measured in the plumage, validating our theoretical approach. This result does not change if we instead model plumage spectra in a bird visual model and measure hue shift as the angular change on the (2-dimensional) hue sphere (see appendix, Figure A.1).

**Figure 4.**
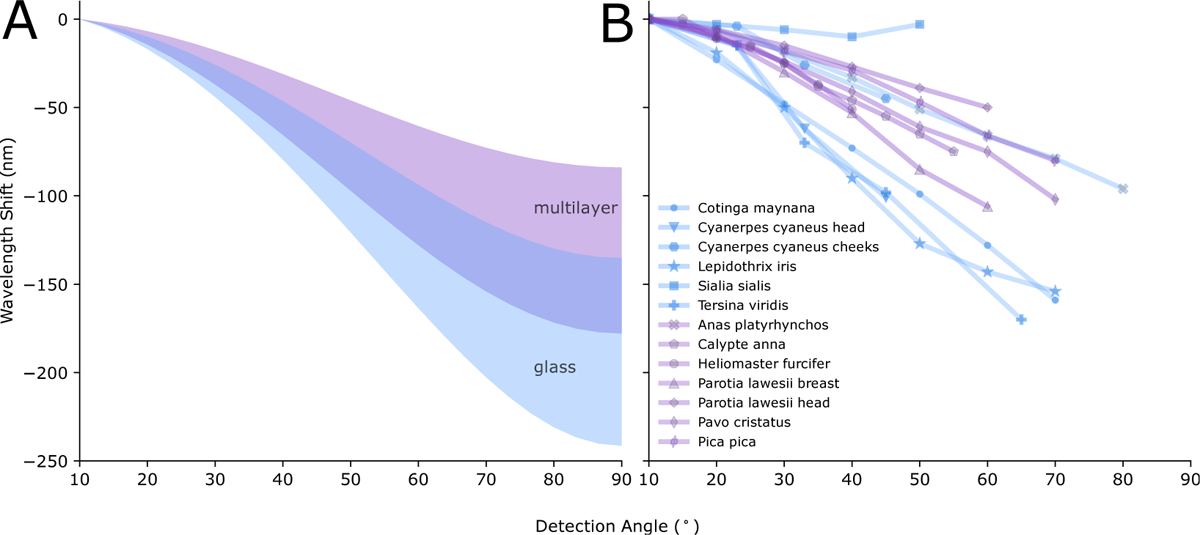
Structural barb (modeled as photonic glass, in blue) and barbule colors (modeled as multilayer, in purple) overlap in iridescence as measured by changing the angle between illumination and detection. Cumulative change in peak spectral wavelength with viewing angle for structural barbules and barbs using A) optical modeling, and B) data from plumage measurements. Data in B) from Stavenga et al. (2017, 2018), Meadows et al. (2011), Freyer Pascal et al. (2019), Gruson et al. (2019), Urquia et al. (2020), Skigin et al. (2019), Noh et al. (2010b), Wilts et al. (2014).

In summary, our results suggest that there is no major difference in either the presence or degree of iridescence between “iridescent” and “non-iridescent” plumage coloration as measured by changing the angle between illumination and detection. However, it is important to point out that photonic glasses are expected to show little to no iridescence when the angle between illumination and detection are kept constant but the sample is rotated (which is theoretically equivalent to rotating the sample under perfectly diffuse lighting)(Noh et al., 2010b, Osorio & Ham, 2002). This is because photonic glasses are rotationally symmetric, while multilayers are not.

## 4. All that glitters is not gold

Why do some objects in nature appear metallic? The natural starting point to answer this question is to explore whether metals have unique reflection properties that distinguish them from other natural objects. Most natural objects reflect light both specularly and diffusely (Figure 5A–B). The diffuse reflection arises from light that has been scattered many times within the material just beneath the surface and exits at a random angle. It can therefore be seen in any direction at equal intensity. The specular reflection arises from the surface has only been scattered once and can therefore only be viewed at the mirror angle. On smooth objects, the specular reflection appears as highlights with sharp edges and is perceived as gloss (Figure 5A). On rough objects, unevenness of the surface will cause specular reflections to spread in multiple directions, mimicking the appearance of a diffuse reflection (Figure 5B). For most natural objects, the diffuse reflection gives rise to the color while the specular reflection has the same spectral distribution as the illuminant and is therefore typically white. In contrast, metals do not follow this typical reflection pattern: they exhibit only specular reflection that is colored independently of the illuminant (Figure 5C–D). Thus, gold has a yellow specular reflection even if the illuminant is white. This unusual reflection property clearly sets metals apart from other natural objects.

**Figure 5.**
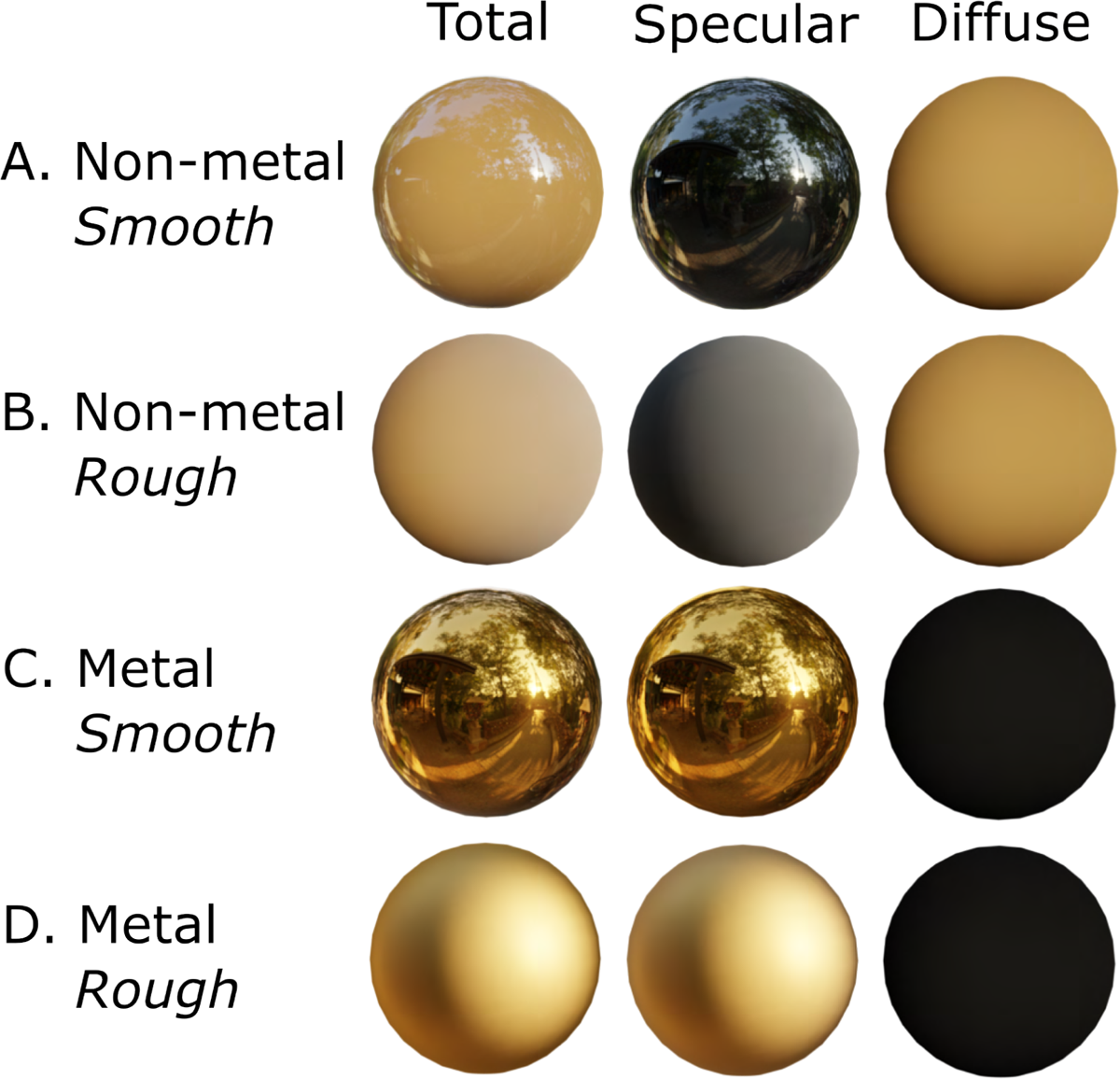
Metals have unique reflection properties: they reflect colored specular light and lack a diffuse reflection. Rendered objects of different materials shown with total reflection, specular reflection only and diffuse reflection only (from left to right). A non-metal object reflects both diffusely and specularly, where the diffuse part confers the color while the specular part has the same spectral distribution as the illumination (A-B). The specular reflection of smooth non-metal objects appear as gloss (A), while on rough objects it appears similar to the diffuse reflection (B). Note, however, that the specular reflection has the same spectral distribution as the illuminant (here a 3D map of a daylight scene in Paris, with blue/white sky and darker ground/vegetation), while the diffuse reflection is colored. Metal objects reflect only specularly (C-D). A rough metal object (C) appears less glossy than a smooth metal object (D). Rendered objects created in Blender v. 3.4.1 (Community, 2022).

In contrast to man-made metallic paints, which simply contain metal particles coupled with a pigment (Rump et al., 2008), structural colors in nature that appear metallic do not contain any metals—they are built with organic materials such as keratin, chitin and melanin. Why do they also appear metallic? If we contrast the reflection properties of structures that appear metallic (structural barbule coloration, Figure 2A), to those that do not (structural barb coloration, Figure 2B), it becomes clear that the structures that appear metallic mimic the reflection pattern of metals. We previously explained that the nanostructures in structural barbules can be approximated by a multilayer structure, while those in structural barbs can be approximated by a photonic glass. Multilayers function as a stack of mirrors, where the specular reflection from each mirror interferes to form a saturated color. Thus, multilayers produce colored specular reflections, just like metals. Moreover, the multilayers are built with highly absorbing melanin pigment (melanosomes, Figure 2A), which nearly eliminates diffuse reflectance. Thus, structural barbule colors incorporate both of the key reflectance properties of metals—a colored specular reflection and very low diffuse reflection. Photonic glasses also produce color from interference effects, but, in contrast to multilayers, they reflect both specular and diffuse light. The specular reflection (in our terminology) is equivalent to what has previously been described as the single-scattered peak (Noh et al., 2010b,c,a, Hwang et al., 2020, 2021) and is colored. However, because photonic glasses also reflect diffusely (multiple-scattered peak), structural barb colors do not meet the dual criteria for metallic appearance (colored specular reflectance *and* low diffuse reflectance).

We are not the first to suggest that metallic luster can be characterized by some combination of reflection properties. In fact, a similar argument was made by Bancroft & Allen (1924) in a paper dedicated to the exploration of “metallic luster”. The authors mentioned two key aspects that are important for the perception of metallic luster: a strong surface reflection and high opacity of the object. Rephrased, this could be understood as equivalent to our condition of a high specular and low diffuse reflection, since “surface reflection” in this context has a similar meaning to our definition of specular reflection, and an opaque body per our definition lacks diffuse reflection (as it does not transmit light and thus no sub-surface scattering can occur). Bancroft & Allen (1924) also suggests that “selective reflection at the surface” gives rise to metallic luster—which is the equivalent of what we have described as a colored specular reflection. We have adopted the term “metallic luster” in this paper to highlight the link to these ideas, albeit having arrived upon them independently.

We suspect that part of the reason that metallic luster has not received more attention in the animal coloration literature is because there is no easy way to quantify it. We therefore propose cross-polarization photography as a method to measure the two key aspects of metallic luster that we have identified (colored specular reflectance and low diffuse reflectance), and we demonstrate this method on a sample of plumage patches colored by diverse mechanisms.

## 5. Measuring metallic luster

With cross-polarization photography, it is possible to separate the specular and diffuse reflection of an object. We therefore used this method to quantify the two key aspects of metallic luster that we have identified above: 1) a colored specular reflection; and 2) a very low diffuse reflection. We measured the first aspect as the saturation of the specular reflection, and the second as the relative specular reflectance (out of the total reflectance, which is the sum of specular and diffuse reflection). A high relative specular reflectance thus equals a low diffuse reflectance.

### 5.1. Plumage selection

We photographed 103 plumage patches from bird specimens at the Princeton Bird Collection (Princeton, New Jersey, USA). We selected species with color mechanisms in the following categories: pigmentary (total of 15 patches, including carotenoid, turacoverdin, psittacofulvin and melanin), structural barb (total of 11 patches), white (which can be considered as a type of structural coloration (Igic et al., 2018); total of 3 patches) and structural barbule (total of 74 patches). The categorization was based on published sources (*n* = 94), and in some cases visual assessment by K. Nordén (*n* = 9, see Appendix C).

### 5.2. Cross-polarization photography

To measure the specular and diffuse reflection from bird plumage, we used cross-polarization photography. This technique takes advantage of the fact that light retains its original polarization when it is single-scattered, but not when it is multiple-scattered. By illuminating the plumage with polarized light, the single-scattered and multiple-scatted part of the reflection can therefore be separated with a second, rotatable polarization filter in front of the camera. Since specular reflections arise from single-scattered light, and diffuse reflections arise from multiple-scattered light, this separates specular and diffuse reflections. Polarization measurements using spectrophotometry have been used previously to study the reflectance properties of photonic structures, including structural barb and barbule coloration (Hwang et al., 2020, Noh et al., 2010b,a,c, Wilts et al., 2014).

Our set-up (Figure 6) consisted of two LED photography lamps which were covered with a polarizing film with a vertical polarization axis. The lamps were set at an approximately 45*^◦^* angle. We used a Nikon D7000 camera, which we mounted on a copy stand and equipped with a rotatable linear polarizer. For each specimen, we took images in the following configurations: 1) with the polarization axis of the camera filter oriented vertically and therefore aligned with the lamps (plane-polarized, PPL, Figure 6A), and 2) with the polarization axis of the camera filter oriented horizontally and therefore perpendicular to the lamps (cross-polarized, XPL, Figure 6B). Using these two sets of images, the specular and diffuse reflectance components of the image can be calculated (see §5.4, *Analysis*). We repeated this process for each patch for a total of three times; each time, we moved the specimen slightly to image a different part of the same patch. The three samples (two images per sample) per patch were collected to get a more representative average of each patch. In each image, we included a color calibration chart (Colorchecker Classic Nano, Calibrate), which allowed us to calibrate camera measurements to known XYZ values (CIE 1931 color-matching functions). All images were captured in RAW format.

**Figure 6.**
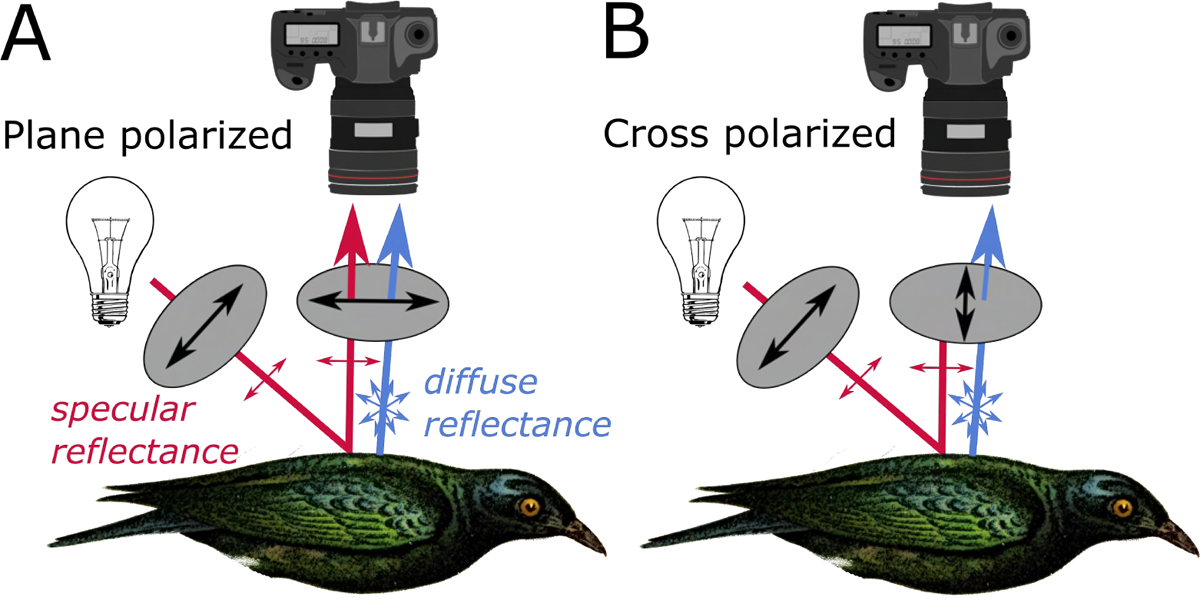
Set-up for cross-polarization photography of bird plumage. The camera was fixed on a copy stand above the specimen, and the lights were fixed at approximately 45*^◦^*. Both camera and light were equipped with linear polarizing sheets. In the plane polarized condition (A), the camera polarizer was set at the same polarizing axis as the lamp, and hence both specular and diffuse reflectance were captured. In the cross-polarized condition (B), the camera polarizer was perpendicular to the axis of the polarizing axis of the lamp, thus only letting through diffuse reflectance.

### 5.3. Image processing and color calibration

We used a custom-written MAT-LAB script to process our RAW images, following Akkaynak et al. (2014) and Sumner (2014). First, images were linearized and demosaiced using bilinear interpolation. We aligned PPL and XPL images using the MATLAB function “imregcorr” to correct for slight shifts between images.

To color calibrate the images, we first combined the plane-polarized image (*I*_ppl_) and the cross-polarized image (*I*_xpl_) for each specimen to produce an image representing the total reflection (*I*_total_). This was necessary because the values of the color standard are based on the total reflection, not the diffuse and specular reflection separately.

We then calculated the linear transformation to map color chart values in *I*_total_ to the known standard XYZ values (CIE 1931 color-matching functions, published for Calibrite Classic Color Checker Nano). This is solved as a linear regression problem of the form:

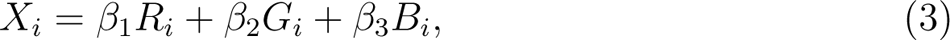

where *R_i_, G_i_* and *B_i_* denote the R, G, and B values respectively of the *i*-th color checker chip in the image *I*_total_, and *X_i_* denotes the standard X, Y, or Z value for the *i*-th color checker chip. The transformation was applied to the images *I*_ppl_ and *I*_xpl_ separately to achieve a color calibrated result. We checked final calibrated values in the image against standard XYZ values to ensure a good fit (see appendix, Figure B.1).

Finally, we converted our XYZ images to RGB space to ease further analysis and interpretation. We converted images to Wide Gamut RGB space, to avoid extensive clipping of values.

### 5.4. Analysis

We selected a patch (700*×*700 px) in each image which represented the patch we wanted to capture, for which all calculations were performed. We calculated the mean intensity values in each color channel for *I*_ppl_ and *I*_xpl_, which we will call *M*_xpl_ and *M*_ppl_, respectively. We chose this approach instead of performing calculations pixel by pixel on the whole patch because it produced more reproducible results. Shiny dark objects are notoriously hard to photograph because correctly exposed highlights will clip values in dark areas of the image and vice versa. Thus, darker areas in our images were more likely to be noisy and out of the calibrated range. Moreover, small shifts in absolute values can significantly shift calculations of ratios (i.e. relative amount of specular reflection and saturation). By averaging intensity over all pixels, we stabilized estimates of dark patches.

#### 5.4.1. Measuring diffuse reflection

The PPL and XPL images can be related to the specular and diffuse reflection as follows:

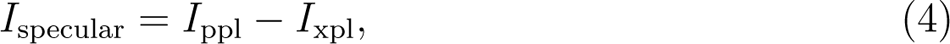

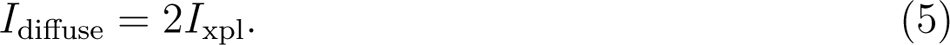

The diffuse reflectance equals twice *I*_xpl_, since a polarization filter will filter out half the intensity of unpolarized light, according to Malus’ law. Thus, from the average intensity values we extracted the mean total intensity (*M*_total_ = *M*_xpl_ + *M*_ppl_), mean specular reflection (*M*_specular_ = *M*_ppl_ *− M*_xpl_) and mean diffuse reflection (*M*_diffuse_ = 2 *× M*_xpl_) for each color channel. Relative specular reflection (*S*) was calculated for the color channel of greatest mean total intensity as follows:

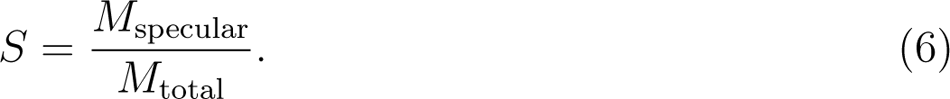

We selected the color channel of greatest mean total intensity, since this would best reflect the plumage color of interest. Thus, we calculated relative specular reflection in the blue color channel for a blue patch, but in the red channel for a red patch.

#### 5.4.2. Measuring colored specular reflection

To measure saturation of the specular reflection, we represented samples in a trigonal RGB color space, i.e. a point (*R, B, G*) *∈* R*_≥_*_0_ gets mapped to

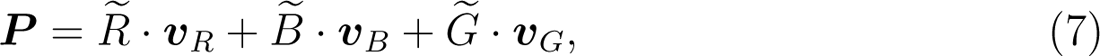

where *R* = *R/*(*R* + *B* + *G*), *B* = *B/*(*R* + *B* + *G*), and *G* = *G/*(*R* + *B* + *G*) are the brightness normalized values and ***v****_R_*, ***v****_B_*, and ***v****_G_* are the vertices of an equilateral triangle. We assume the triangle to be centered at the origin, i.e. ***v****_R_* + ***v****_B_* + ***v****_G_* = **0**, such that the unsaturated color (white) is mapped to the origin **0**. Saturation, in this space, is then represented as the ratio of the distance of the point ***P*** from the origin and the length of the line segment of a ray from the origin through the point ***P*** contained within the triangle. Saturation therefore represents the relative imbalance of the values *R, B, G* (see Equation (8)). Explicitly, saturation Sat(***P***) of the point ***P***, given by Equation (7), is given by

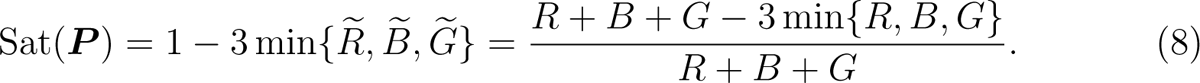

This may be seen as follows. For ***P*** = **0**, we have Sat(**0**) = 0 and 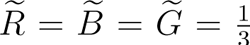. For ***P*** *̸*= **0**, we may suppose without loss of generality that 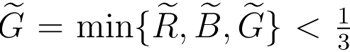. Then, Sat(***P***)*^−^*^1^ is equal to the scalar *t* such that *t ·* ***P*** lies on the edge *E_RB_* between ***v****_R_* and ***v****_B_*. We have

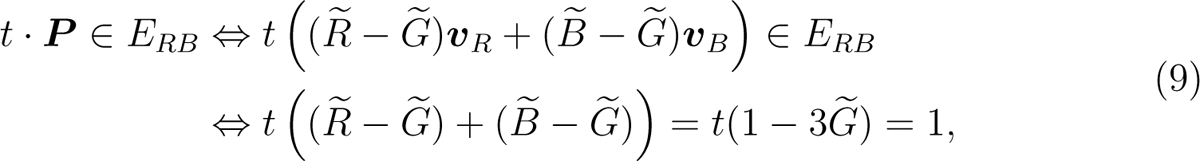

where we have used ***v****_R_*+ ***v****_B_*+ ***v****_G_*= **0** and *α****v****_R_* + *β****v****_B_ ∈ E_RB_ ⇔ α* + *β* = 1 and *α, β ≥* 0.

### 5.5. Results

In Figure 7A, the results of the cross-polarization photography are visualized in terms of specular saturation and relative specular reflection of each patch. Of the plumage color types we tested, structural barbule colors stand out as the only type that has a high relative specular reflection that is saturated (Figure 7A–B). They vary in the magnitude of specular saturation, which is likely tied to the number of melanosome layers in the nanostructure (Figure 2A). As the number of layers increase in a multilayer, spectral peak height increases while spectral peak width decreases (Kinoshita et al., 2008). Patches that record high specular saturation will therefore appear very bright and saturated (e.g. Resplendent quetzal (*Pharomachrus mocinno*), Figure 2Aa) while patches that record low specular saturation will appear less bright and saturated (e.g. Common raven (*Corvus corax*), Figure 2Ab). Thus, cross-polarization photography successfully captures the unique reflectance properties of plumage with metallic luster—proving that this could be used as a proxy to quantify it.

**Figure 7.**
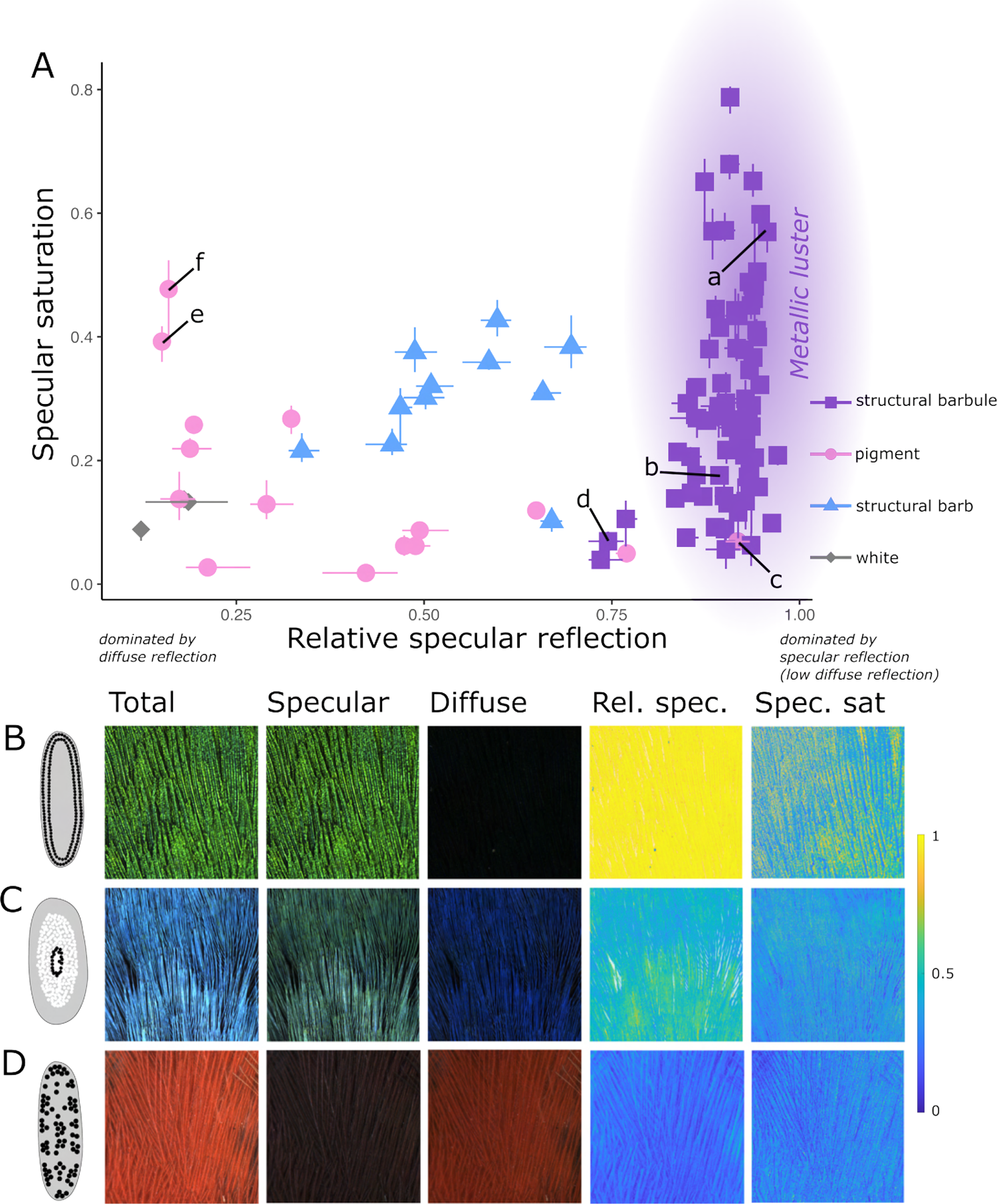
Cross-polarization photography effectively separates plumage patches with metallic luster (structural barbules) from other types of plumage. A, Relative specular reflection and specular saturation of plumage patches with different color mechanisms. Data points represent the average measurement of three samples (per patch), the lines represent the range of the samples. Lower case letters in A mark specific patches discussed in the result: a, *Pharomachrus mocinno*; b, *Corvus corax*; c, *Gallus gallus*; d, *Picoides pubescens*; e, *Piranga olivacea*; f, *Carduelis tristis*. B-D, example patches of each type of plumage shown as a total intensity image, specular image, diffuse image, and false-color images of relative specular reflection and specular saturation. B, structural barbule (Resplendent quetzal, (*Pharomachrus mocinno*)); C, structural barb (Swallow tanager, (*Tersina viridis*)); D, pigmentary (Northern cardinal, (*Cardinalis cardinalis*)).

Plumage produced with color mechanisms other than structural barbules occupy different areas of the graph (Figure 7A). Structural barb colors cluster in the middle of the graph, suggesting that they reflect both diffuse light and colored specular light— in line with our expectations (Figure 7A, C). Structural white, on the other hand, exhibits low relative specular reflection, due to its very high diffuse reflectance (Figure 7A). Similarly, pigmentary patches primarily record low relative specular reflection and saturation (Figure 7A, D). However, there are two interesting exceptions to these general patterns.

Firstly, some pigmentary patches show high proportions of specular reflection and therefore plot in the lower right quadrant, close to patches with structural barbule coloration. These high specular components are present for pigmentary patches that are very dark (melanin based) yet glossy (e.g. Junglefowl (*Gallus gallus*), Figure 7Ac). The difference between dark glossy patches and patches with structural barbule coloration is that the gloss is unsaturated (white) for the former and colored for the latter. However, the distinction between white and colored is a nuanced question. Quantifying this difference well requires very precise measurements that are beyond the capabilities of the digital photography we used here. Thus, patches with structural barbule coloration that produce very low specular saturation overlap with dark pigmentary and glossy patches in the lower right quadrant, e.g. Downy woodpecker (*Picoides pubescens*), Figure 7Ad. Nevertheless, this overlap represents a real gradual transition between glossy pigmentary plumage and plumage with metallic luster that is mirrored on the nanostructural level in the feather. Structural barbules and melanin pigmented barbules both consist of keratin and melanin, only differing in the extent to which the melanosomes are ordered (Figure 2A, C). In general, higher order of the melanosomes increases the saturation of the specular reflection. Some patches classified as structural barbule coloration with low specular saturation exist in this transition zone between glossy pigmentary plumage and metallic luster.

Secondly, some pigmentary patches record a saturated specular reflection in combination with a diffuse reflection (e.g. Scarlet tanager (*Piranga olivacea*), Figure 7Ae and American goldfinch (*Carduelis tristis*), Figure 7Af). This is surprising, because pigments typically produce color through the diffuse reflection, and we would expect the specular part to be white (Figure 5A). Intriguingly, the patch which records the most saturated specular reflectance is the yellow back of the American goldfinch (*Carduelis tristis*, Figure 7Af)—a species which has previously been shown to amplify pigmentary color using feather nanostructures (Shawkey & Hill, 2005). Air-filled vacuoles in the barbs function as mirrors reflecting light behind a carotenoid pigmented keratin layer. Thus, the specular colored light recorded for some pigmentary patches may in fact signal a structural amplification of the pigmentary color. This would need to be confirmed with additional studies of feather nanostructures—but if our inference is correct our results suggests structural amplification of carotenoid colors may be widespread in birds.

In summary, our results show that cross-polarization can be used to quantify a proxy of metallic luster, by measuring characteristics of the specular reflection.

## 6. Discussion

We have argued that the term “iridescence” is often used in animal coloration literature to describe metallic luster, and that this is a separate property from iridescence as commonly defined (a change in peak spectral wavelength with viewing or illumination angle). We quantify metallic luster using two reflection characteristics (a colored specular reflection, measured by specular saturation and a low diffuse reflection, measured by relative specular reflectance), and show using bird plumage that while “iridescent” (structural barbule) and “non-iridescent” (structural barb) plumage both exhibit iridescence when changing the angle between illumination and observation (Figure 4), the latter lacks metallic luster (Figure 7). In other words, structural barbule colors exhibit both iridescence and metallic luster, while structural barb colors exhibit iridescence but lack metallic luster. By separating iridescence and metallic luster, we clarify the terminology surrounding structural colors and construct a framework that can be applied to any structural coloration in nature, not only bird coloration.

In this section, we would like to explore two general critiques that may be raised in response to introducing metallic luster as a property of animal coloration. Firstly, while we might be able to measure this property, does it have any biological significance? For example, does the presence or absence of metallic luster in a signal matter to the receiver? Secondly, even if metallic luster has biological significance, how good is the measure we have proposed at capturing metallic luster, which in reality is a perceptual rather than an objective quality?

### 6.1. Biological significance of metallic luster

Objects with metallic luster appear to stand out to human observers: it is no coincidence that beetles and butterflies with metallic luster are favorite objects of collectors (Finet, 2023), and bird feathers with metallic luster have been used to adorn clothing ranging from Aztec headdresses (McMahon, 2017) to ladies hats in the 1800s (Eluwawalage, 2015). But is there any reason to assume that metallic luster is a salient feature to other animals? If not, one might argue that even though it is a measurable property that matters to humans, it has little biological significance and no role to play in the evolution of these colors.

There are not yet any studies investigating directly whether animals pay attention to objects with metallic luster—however, we can make a plausible argument that they do, based on the shared evolutionary role of vision in animals. For humans, the metallic luster of an object immediately tells us something about its material properties. It is therefore part of what has been termed our material perception, or our ability to estimate complex material properties from visual cues (e.g. surface texture, density, and viscosity (Fleming, 2017)). Material perception is critical for our ability to interact appropriately with objects in our environment—for example it allows us to predict the weight of an object or whether a surface is wet or dry. Gloss in particular has been found to be of great importance for material perception (Fleming, 2017, Schmid et al., 2023, Ged et al., 2010). By adjusting specular strength, specular saturation and the smoothness of computer-rendered objects shown on a screen, Schmid et al. (2023) found that people categorized objects into distinct categories such as “porcelain”, “wax”, “metal” and “plastic”.

Classifying materials such as “plastic” or “porcelain” may not be relevant to most animals—but predicting the material properties of an object certainly is. Just like object detection probably has deep evolutionary roots (Soto & Wasserman, 2014, Jitsumori & Delius, 2001, Schumacher et al., 2016), material perception is likely widespread among other animals. For animals that rely on vision, cues derived from specular reflections are likely to be important for this task, since the appearance of the specular reflection is tightly linked to material properties. Moreover, there is limited direct evidence for gloss perception in non-human animals—Okazawa et al. (2012) recorded specific neural responses to viewing glossy surfaces in macaques.

Thus, based on the key role of gloss perception for object and material recognition in humans, it is likely that many animals (that rely on vision) also perceive it (Franklin & Ospina-Rozo, 2021). Since an important feature of metallic luster is colored specular reflection (or colored gloss), it is likely that many animals would also perceive metallic luster. This could be tested using behavioral trials.

However, there is a difference between being able to detect metallic luster and being attracted to it. Humans tend to prefer shiny or glossy objects, including objects with metallic luster, over matte objects (Silvia et al., 2018, Meert et al., 2014, Silvia et al., 2021). The reason for this preference is still unknown (Fleming, 2017). Coss (1990) suggested that the attraction to glossy surfaces is an evolved adaptation to find water in terrestrial environments. He supported his argument by an experiment in which people rated papers with glossier surfaces as “wetter” than papers with matte surfaces (Coss, 1990). Coss et al. (2003) also conducted an experiment which showed that toddlers tended to lick glossy or metallic plates significantly more than matte plates— a behavior they interpreted as an indication that the toddlers perceived the glossy surfaces as wet. Alternatively, glossy or lustrous surfaces may capture our attention because they register as a movement (Braun & Braun, 1995). Visual perception of motion is critical to a variety of tasks, including judging the shape, depth, movement, and speed of objects, as well as identifying other animals in our environment (Sekuler et al., 2002). Since movement of objects catches our attention (Franconeri & Simons, 2005, 2003, Abrams & Christ, 2003), it is possible that glossy or lustrous surfaces do the same by mimicking or amplifying movement cues. Surfaces with metallic luster may additionally be attractive because they have unusual reflective properties—while many objects are glossy, only a few have a colored shine.

Little is known about animal preference or attention to glossy objects (Franklin & Ospina-Rozo, 2021). A few studies have explored preference for shiny objects in corvids, who according to widely held beliefs in European folklore collect shiny objects. Behavioral trials do not support this, showing only that corvids have a preference for exploring novel objects (Shephard et al., 2015, Heinrich, 1995, Jacobs et al., 2014). In addition, a couple of studies have investigated how glossiness affects the detectability of prey. Specifically, these studies compared the attack rates (by bird predators) in the wild on replica beetles with either glossy or matte elytra. In one study, survival rates between glossy and matte beetles of a similar green hue placed in natural environments did not differ (Franklin et al., 2022), while a more recent study found that glossy beetles of three different hues were consistently easier to detect for bird predators than matte replicas (Thomas et al., 2014). Thus, while there is not yet much direct evidence to suggest that animals pay attention to glossy objects, it has not been tested in many species or contexts. The attention to or preference for metallic luster specifically has, to our knowledge, never been investigated in animals.

Despite the scarcity of evidence, there is reason to believe that metallic luster may have wider biological significance since the two proposed hypotheses for the human preference for glossy surfaces—ability to identify water and to perceive motion—are likely to be important to a wide range of animals beyond humans. This hypothesis is particularly intriguing in birds, which have good vision and are known to incorporate metallic luster in plumage patches used for display. Previous studies have focused on how brightness, hue, saturation and in some instances iridescence of such plumage patches influence mate choice or are tied to animal condition (Bitton et al., 2007, Loyau et al., 2007, Dakin & Montgomerie, 2013, Simpson & McGraw, 2019, Hill et al., 2005). For example, brighter plumage has been linked with increased mating success in Peafowl (*Pavo cristatus*, (Loyau et al., 2007)) and Tree swallow (*Tachycineta bicolor*, (Bitton et al., 2007)). Introducing metallic luster as a property of coloration allows us to ask whether it is the brightness in general that is important (which could arise from either diffuse or specular reflections) or the presence of metallic luster (bright and colored *specular* reflections only). On a macroevolutionary level, we could investigate what kind of environments or signal types may favor the evolution of metallic luster (as opposed to diffuse coloration such as pigmentary traits).

Thus, we would argue that there are many interesting biological questions that could be asked using the concept of metallic luster. Of course, to test these hypotheses we need a way to quantify it. Can the measure of metallic luster we developed in this paper (i.e., using cross-polarization photography to identify colors with a colored specular reflection and low diffuse reflection) really capture this property, which as we have seen likely involves higher level visual processing?

### 6.2. Metallic luster as a Gestalt-like property

Our measure of metallic luster focuses on particular reflective properties of surfaces. Yet, it is clear that metallic luster must be a perceptual rather than objective property of objects, because it is known that individuals can sometimes differ in their judgment of whether an object has metallic luster (Bancroft & Allen, 1924, Todd & Norman, 2018). This is why we have called our measure a proxy, just like a shift in peak spectral wavelength can be considered a proxy of the perceptual experience of iridescence. Proxies are useful since they allow us to quantify a property that is hard to measure directly—but by virtue of being proxies they can also be misleading in some contexts. Bancroft & Allen (1924) point out that variations in light intensity on a surface either in space or time is critical to the perception of metallic luster. A perfectly smooth mirror does not look metallic—it is only when some unevenness in the surface is introduced that it appears metallic. Our definition and measurement of metallic luster cannot distinguish between these two situations. Neither can our measure distinguish between a surface that is silvery and one that is dark and glossy (both have high specular reflectance but lack specular saturation). In fact, even human judgments of whether an objects is black and glossy or silvery can in some situations be highly variable (Todd & Norman, 2018).

Evidence emerging from studies of material perception and gloss is supporting the idea that the perception of gloss and metallic luster is dependent on a variety of cues, including the location and shape of specular highlights, their movement across the object, the contrast in highlights and low-lights, the surface texture of an object as well as the illumination characteristics (Chadwick & Kentridge, 2015, Schmid et al., 2023, Todd & Norman, 2018, Norman et al., 2020). Moreover, the expectation of an object being of a particular material affects how we perceive it. Matsumoto et al. (2015) found that objects that had a similar hue to gold were perceived as more glossy and metallic than objects that did not, despite having the same reflective properties. In another experiment, it was found that objects that feel smooth to the touch are perceived as more glossy than rough objects (Adams et al., 2016). Our prior knowledge and expectation of an object or material can thus also directly influence how we perceive it (Alley et al., 2020).

These insights from studies of human perception of gloss and metallic luster suggest it can probably be best understood as a gestalt-like property (Schmid et al., 2023). Gestalt psychology originated in the early 1900s (Wagemans et al., 2012), but it has recently received renewed attention in visual perception research (Jäkel et al., 2016). A core concept of Gestalt theory is that the whole can be more than the sum of its parts (superadditivity) (Jäkel et al., 2016, Wagemans et al., 2012). To state this a bit more concretely in relation to metallic luster, we can see that there is no single point, or combination of points, in a color space, that can be called “gold”. Rather, it is the particular distribution of colors and highlights of an object (the Gestalt) that together create the appearance of gold (Figure 8, cf. Komatsu et al., 2013).

**Figure 8.**
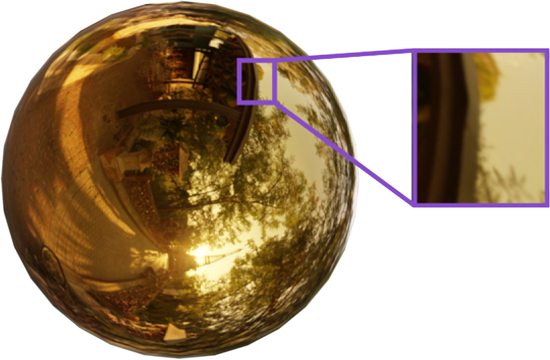
Metallic luster is probably a gestalt-like property that arises from the combination of many different cues, including the interaction between highlights and object shape. The cut-out of the sphere does not look golden on its own—it is only as part of the sphere as a whole that it appears golden. Figure based on Komatsu et al., 2013.

Gestalt psychology is not new to the field of animal coloration—but it has hitherto been limited to the study of animal camouflage, which was a favorite example of early Gestalt theorists (Osorio & Cuthill, 2015). However, a major problem with Gestalt psychology is that it is often hard to precisely quantify the concepts (Wagemans et al., 2012). We do not offer here any solution to this problem, but we argue that the Gestalt is a useful way of thinking of properties like metallic luster, gloss and iridescence—even if we cannot yet quantify it as a Gestalt. That is to say, even though measurements of metallic luster or iridescence using proxies based on separate cues are useful, we need not assume that the perception itself operates in a similar way. For example, if we view iridescence as the combination of many different hues, each seen at a specific angle of light and observation, it appears to be a very complicated signal to remember (Stuart-Fox et al., 2020, Thomas et al., 2023). If we instead consider iridescence as a holistic Gestalt, we need not assume that this signal is more complicated or harder to remember than any other.

In summary, the measure of metallic luster—a colored specular reflection in combination with low diffuse reflectance—we have proposed in this paper should be viewed as a proxy, rather than the equivalent, of metallic luster. This measure could be improved by adding additional features (e.g. contrast and coverage of high- and low-lights in the image)—although this is perhaps best done in step with increasing knowledge of different animals’ perception of luster. Fundamentally, metallic luster is probably a gestalt-like quality rather than a single metric. Nevertheless, in the absence of a way to measure metallic luster directly, a simple proxy can still be a useful starting point to explore basic questions about this understudied aspect of animal coloration.

## 7. Conclusion

That some objects in nature appear metallic has probably been appreciated by humans for as long as they have encountered animals and minerals with structural colors. Therefore, our goal in this paper is not to present the new idea that some natural objects have metallic-like reflections but to highlight and develop a concept that has long existed, albeit in disparate literatures and with little direct examination. In particular, we argue that iridescence and metallic luster have become blurred in the literature and we show using examples and data that they are in fact two distinct properties that may or may not occur together.

We developed a measure of metallic luster which is based on the unique reflection properties of metals, which successfully quantifies metallic luster in bird plumage. While this method of measuring metallic luster is quite simple, it is a useful starting point for exploring hypothesis about the importance of metallic luster to animals signals.

## Acknowledgements.

We would like to thank Elizabeth Horn for allowing us to use the specimens of the Princeton Bird Collection for our research and Roel Muñez for generously lending us a copy stand and offering general advice on photography. R.S. wishes to extend his gratitude to his employer, the Institute for Mathematical Research (FIM) at ETH Zürich; while A.B.S. was supported by funding from the Princeton University Dean for Research, the High Meadows Environmental Institute, and a gift from William H. Miller III. We further appreciate funding to M.C.S. from Princeton University, the National Science Foundation (Award 2029538), a Packard Fellowship for Science and Engineering, and the generosity of Eric and Wendy Schmidt by recommendation of the Schmidt Futures Polymaths Program.

## Appendix A. Iridescence as a change in hue

We have defined iridescence as a shift in peak spectral wavelength with viewing or illumination angle—not as a shift in hue with viewing or illumination angle. This is partly out of convenience, because the former is agnostic to the viewer’s visual capabilities, and partly because it is how iridescence is typically measured in the literature.

Would our conclusions differ if we instead measured iridescence as a shift in hue with illumination angle?

Generally, the answer is “no” since peak spectral wavelength is a rough approximation of hue if measured within the visual range of the viewer. However, there are two exceptions to this.

Firstly, spectra that have multiple peaks instead of a single main peak will be poorly described by the peak spectral wavelength. Moreover, if the relative heights of multiple peaks in a spectrum change over varying incidence angles, the peak spectral wavelength might not describe the gradual change of a single peak, but jump between different peaks for each angular measurement. In such a situation, we would not expect hue shift to roughly approximate the shift in peak spectral wavelength.

The second exception is spectra with very broad peaks. Multilayer structures that produce broad peaks (broadband reflectors) give rise to silvery or golden appearances, and can be found in for example some beetles (Seago et al., 2009). The broadband reflection arise from variations in the layer thicknesses in the stack, which results in constructive interference at multiple wavelengths (Seago et al., 2009, Parker et al., 1998). Just like any other multilayer reflector, the spectral peak will shift as the angle of light or observations is changed. However, this effect is hardly perceivable, since the reflectance peak is very broad. Thus, the golden Christmas beetle (*Anoplognathus aureus*) appears to a human observer golden just like the metal gold—despite exhibiting a measurable shift in spectral peak location (Ospina-Rozo et al., 2022). Therefore, such broadband reflectors might be considered iridescent if measured as a shift in peak spectral wavelength (or here spectral location, which captures the same property for broad peaks), but not if measured as a shift in (human) hue.

In §3.2, *Iridescence*, we compare the iridescence of plumage with structural barbule and barb coloration by plotting their cumulative shift in peak spectral wavelength over increasing specular angles. Since the spectra we used had well defined single peaks that were not very broad, we would expect a similar result if we instead had measured hue shift. However, to test this directly we also calculated hue shifts using a bird visual model. We included all species where we had access to the raw spectral data (12 species). Spectra were processed and modeled in a bird visual model using the R package *pavo* (Maia et al., 2013), (cone sensitivities set as “avg.uv” for all species). Using the relative quantum cone catches from the visual model, we mapped each spectrum to coordinates in a tetrahedral color space, where each vertex represents one cone in the avian eye (Endler & Mielke, 2005, Stoddard & Prum, 2008). The hue shift was then calculated as the (cumulative) change in angle between these points (arising from spectra measured at varying specular angles).

The results (Figure A.1) are comparable to the earlier results for peak spectral wavelength (Figure 4B)—structural barbule and structural barb coloration broadly overlap in iridescence. Thus, interpreting iridescence as a hue shift with viewing or observation angle does not change our conclusion.

**Figure A.1.**
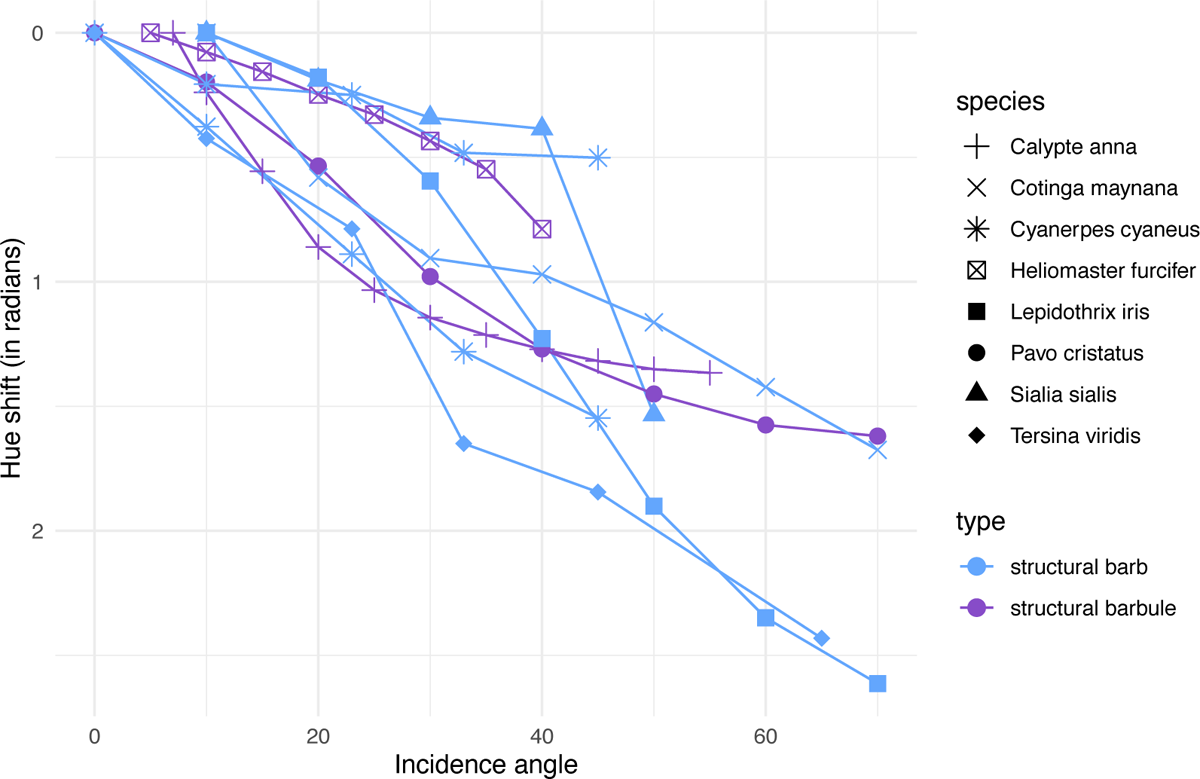
Hue shift over varying angles (specular configuration) for species with structural barbule coloration (purple), and structural barb coloration (blue). Data from Meadows et al. (2011), Freyer Pascal et al. (2019), Gruson et al. (2019), Urquia et al. (2020), Skigin et al. (2019), Noh et al. (2010b).

## Appendix B. Validation of image color calibration

**Figure B.1.**
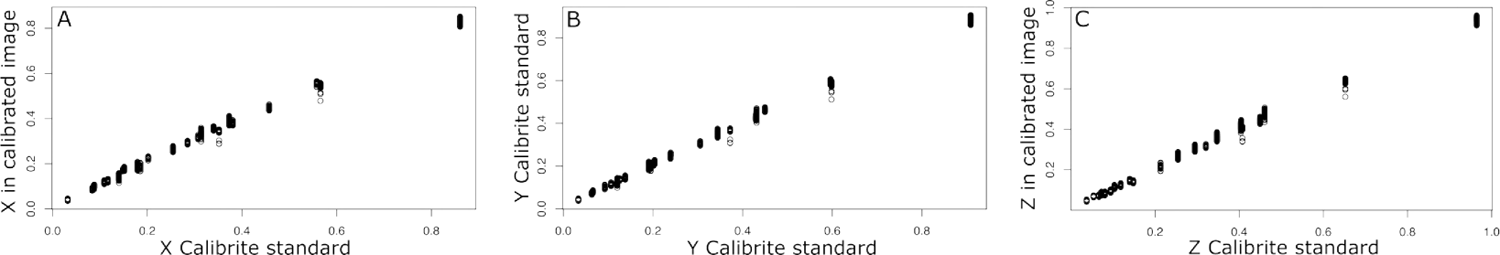
Calibrated XYZ values for each calibrated image plotted against the published Calibrite XYZ standard values, for X (A), Y (B) and Z (C). All images show good convergence to the standard.

## Appendix C. Specimens imaged with cross-polarization photography

**Table C.1.**
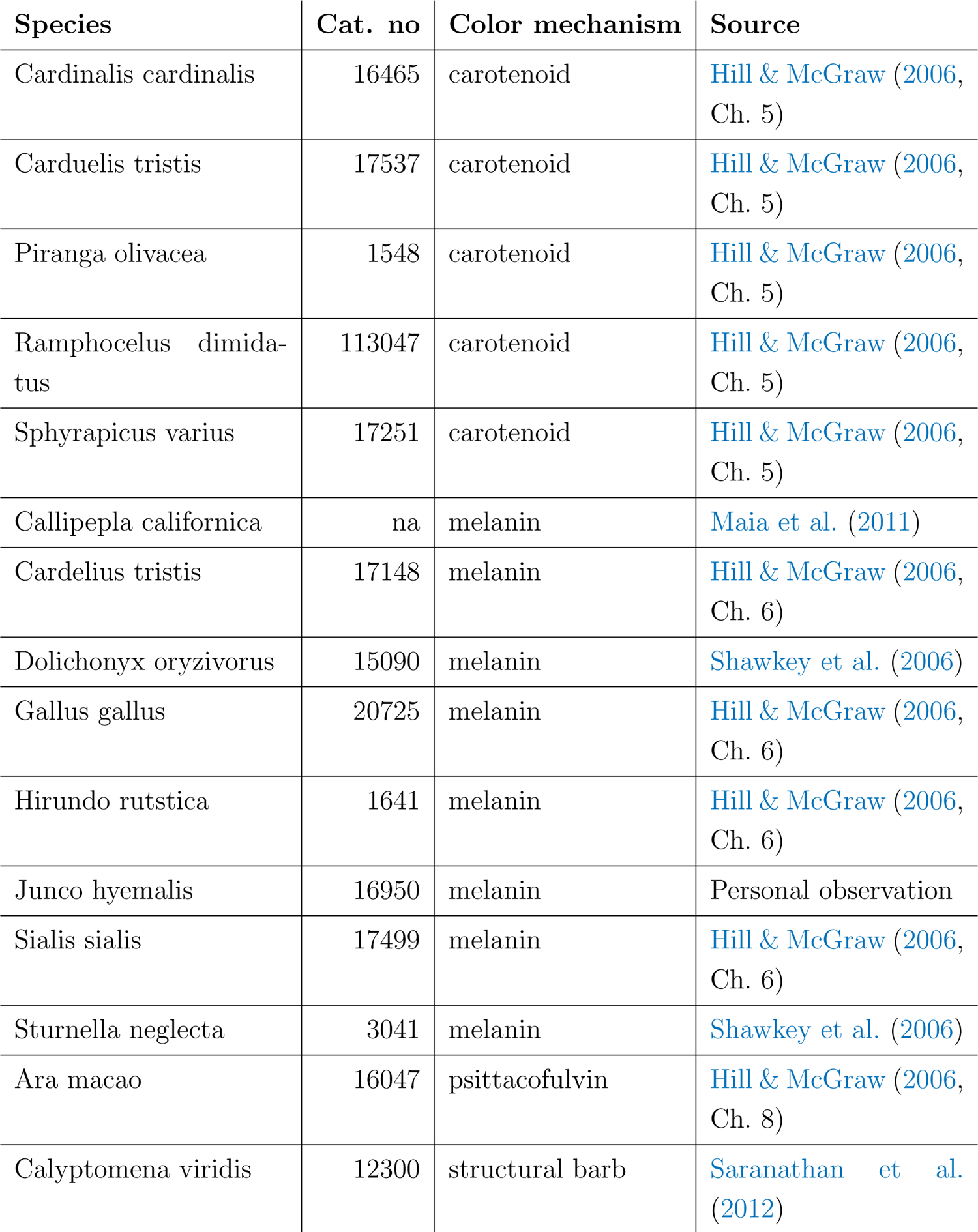
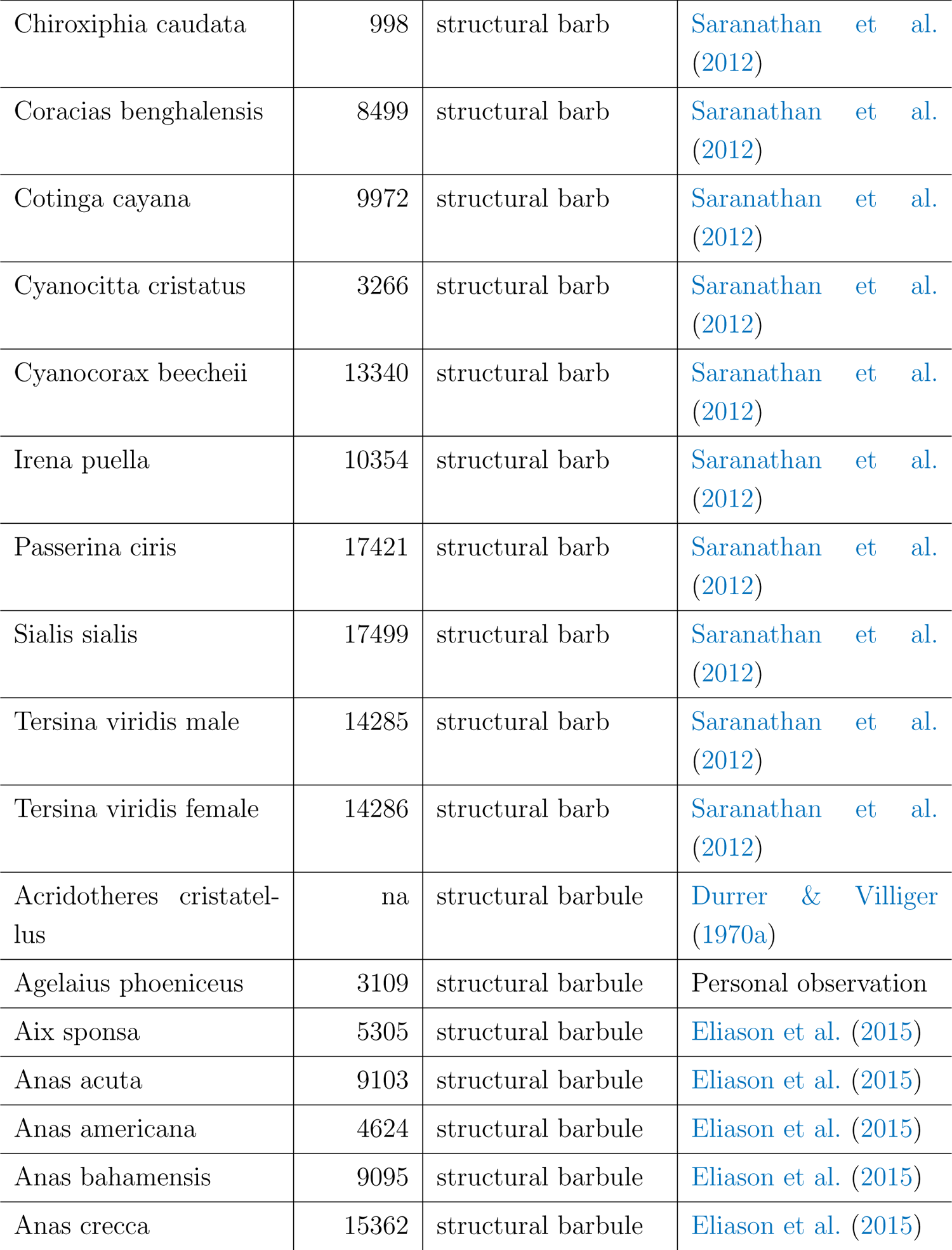
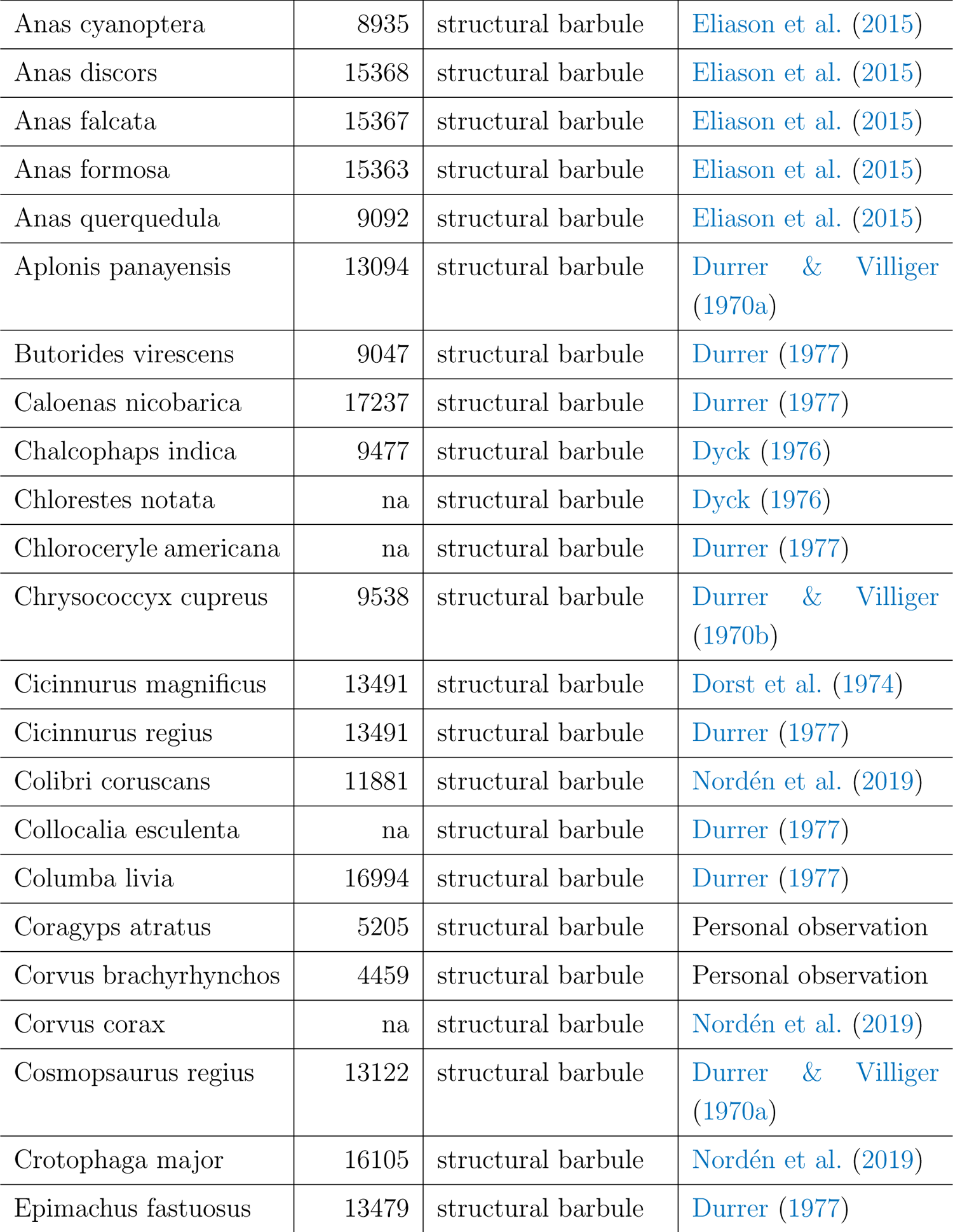
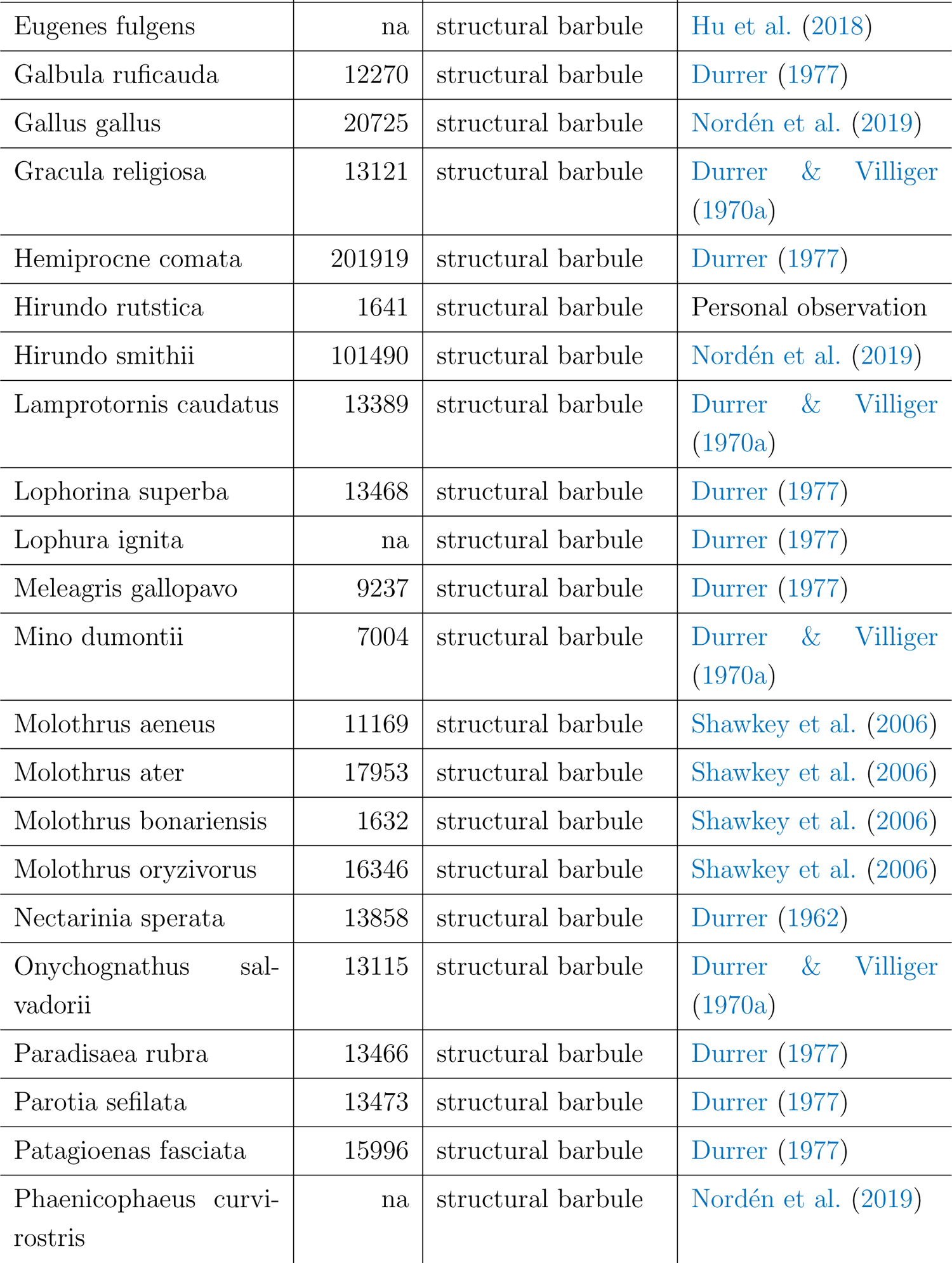
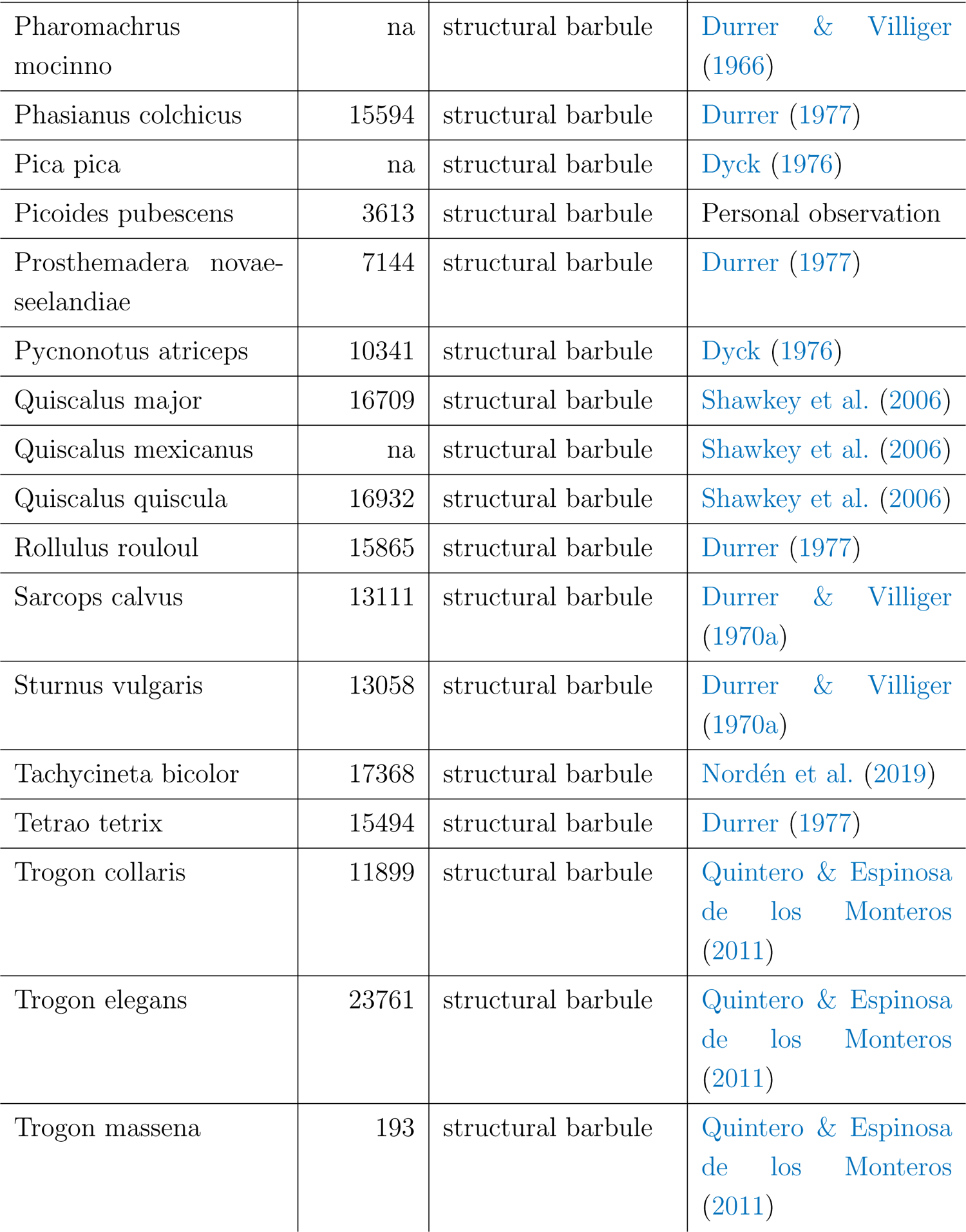
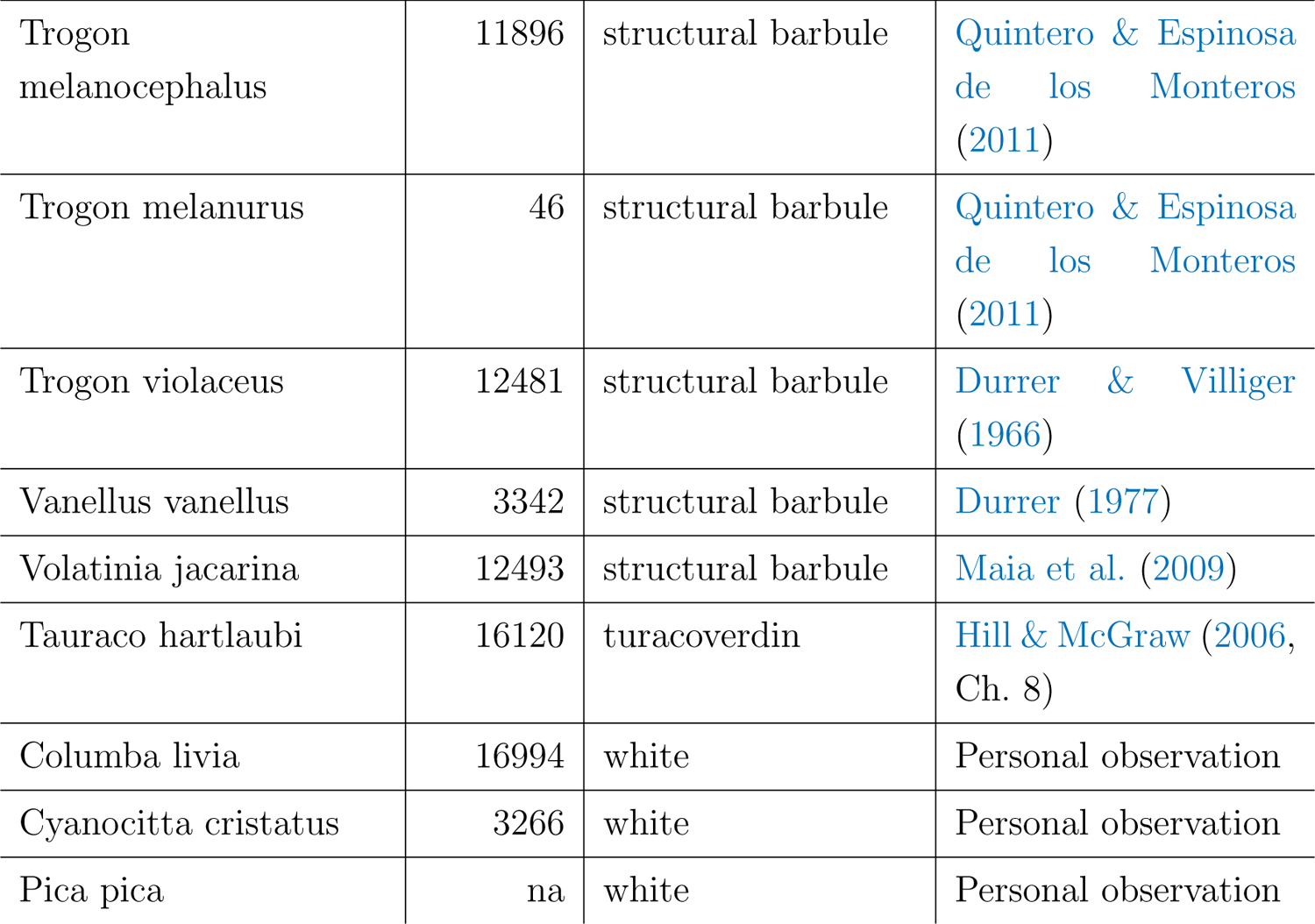
Specimens imaged with cross-polarization photography (all from the Princeton Bird Collection). The reference for each species’ plumage color mechanism is listed under “Source”.

## References

Abrams, R. A. & Christ, S. E. (2003). Motion onset captures attention. Psychological science, 14(5), 427–432.

Adams, W. J., Kerrigan, I. S., & Graf, E. W. (2016). Touch influences perceived gloss. Scientific Reports, 6(1), 21866.

Akkaynak, D., Treibitz, T., Xiao, B., Gurkan, U. A., Allen, J. J., Demirci, U., & Hanlon, R. T. (2014). Use of commercial off-the-shelf digital cameras for scientific data acquisition and scene-specific color calibration. JOSA A, 31(2), 312–321.

Allaby, M., Ed. (2013). A dictionary of geology and earth sciences. Oxford paperback reference. Oxford: Oxford University Press, fourth edition edition.

Alley, L. M., Schmid, A. C., & Doerschner, K. (2020). Expectations affect the perception of material properties. Journal of Vision, 20(12), 1–1.

Auber, L. (1957). The distribution of structural colours and unusual pigments in the class Aves. Ibis, 99(3), 463–476.

Bancroft, W. D. & Allen, R. (1924). Metallic luster. II. The Journal of Physical Chemistry, 29(5), 564–586.

Bitton, P.-P., O’Brien, E. L., & Dawson, R. D. (2007). Plumage brightness and age predict extrapair fertilization success of male tree swallows, Tachycineta bicolor. Animal Behaviour, 74(6), 1777–1784.

Braun, J. H. & Braun, S. W. (1995). Speculations about the value of gloss. Color Research & Application, 20(6), 397–398.

Chadwick, A. C. & Kentridge, R. W. (2015). The perception of gloss: A review. Vision Research, 109, 221–235.

Community, B. O. (2022). Blender - a 3D modelling and rendering package.

Coss, R. G. (1990). All that Glistens: Water Connotations in Surface Finishes. Ecological Psychology, 2(4), 367–380.

Coss, R. G., Ruff, S., & Simms, T. (2003). All That Glistens: II. The Effects of Reflective Surface Finishes on the Mouthing Activity of Infants and Toddlers. Ecological Psychology, 15(3), 197–213.

Dakin, R. & Montgomerie, R. (2013). Eye for an eyespot: how iridescent plumage ocelli influence peacock mating success. Behavioral Ecology, 24(5), 1048–1057.

Dorst, J., Gastaldi, G., Hagege, R., & Jacquemart, J. (1974). Différents aspects des barbules de quelques Paradisaeidés sur coupes en microscopie électronique. Comptes Rendus de l’Academie des Sciences Paris, 278, 285–290.

Doucet, S. M. & Meadows, M. G. (2009). Iridescence: a functional perspective. Journal of The Royal Society Interface, 6, S115–S132.

Durrer, H. (1962). Schillerfarben beim Pfau (Pavo cristatus L.). Verhandlungen der Naturforschenden Gesellschaft in Basel, 73(1), 204–224.

Durrer, H. (1977). Schillerfarben der Vogelfeder als Evolutionsproblem. PhD Thesis, University of Basel.

Durrer, H. & Villiger, W. (1966). Schillerfarben der Trogoniden. Journal fur Ornithologie, 107(1), 1–26.

Durrer, H. & Villiger, W. (1970a). Schillerfarben der Stare (Sturnidae). Journal fur Ornithologie, 111(2), 133–153.

Durrer, H. & Villiger, W. (1970b). Schillerradien des goldkuckucks (Chrysococcyx cupreus (Shaw)) im elektronenmikroskop. Zeitschrift fur Zellforschung und Mikroskopische Anatomie, 109(3), 407–413.

Dyck, J. (1976). Structural colours. Proceedings of the International Ornithological Congress, 16, 426–437.

Eliason, C. M., Maia, R., & Shawkey, M. D. (2015). Modular color evolution facilitated by a complex nanostructure in birds. Evolution, 69(2), 357–367.

Eluwawalage, D. (2015). Exotic fauna and flora: fashion trends in the nineteenth century. *International Journal of Fashion Design*, Technology and Education, 8(3), 243–250.

Endler, J. A. & Mielke, P. W. (2005). Comparing entire colour patterns as birds see them. Biological Journal of the Linnean Society, 86(4), 405–431.

Finet, C. (2023). Light as matter: natural structural colour in art. Humanities and Social Sciences Communications, 10(1), 1–14.

Fleming, R. W. (2017). Material Perception. Annual Review of Vision Science, 3(1), 365–388.

Franconeri, S. L. & Simons, D. J. (2003). Moving and looming stimuli capture attention. Perception & Psychophysics, 65(7), 999–1010.

Franconeri, S. L. & Simons, D. J. (2005). The dynamic events that capture visual attention: A reply to Abrams and Christ (2005). Perception & psychophysics, 67(6), 962–966.

Franklin, A. M. & Ospina-Rozo, L. (2021). Gloss. Current Biology, 31(4), R172–R173.

Franklin, A. M., Rankin, K. J., Ospina Rozo, L., Medina, I., Garcia, J. E., Ng, L., Dong, C., Wang, L.-Y., Aulsebrook, A. E., & Stuart-Fox, D. (2022). Cracks in the mirror hypothesis: High specularity does not reduce detection or predation risk. Functional Ecology, 36(1), 239–248.

Freyer Pascal, Wilts Bodo D., & Stavenga Doekele G. (2019). Reflections on iridescent neck and breast feathers of the peacock, Pavo cristatus. Interface Focus, 9(1), 20180043.

Gadow, H. (1882). On the Colour of Feathers as affected by their Structure. Proceedings of the Zoological Society of London., 50(3), 409 – 422.

Ged, G., Obein, G., Silvestri, Z., Le Rohellec, J., & Vienot, F. (2010). Recognizing real materials from their glossy appearance. Journal of Vision, 10(9), 18–18.

Ginneken, B. v., Stavridi, M., & Koenderink, J. J. (1998). Diffuse and Specular Reflectance from Rough Surfaces. Appl. Opt., 37(1), 130–139.

Gruson, H., Andraud, C., Daney de Marcillac, W., Berthier, S., Elias, M., & Gomez, D. (2019). Quantitative characterization of iridescent colours in biological studies: a novel method using optical theory. Interface Focus, 9(1), 20180049.

Haecker, V. (1890). Ueber die Farben der Vogelfedern. Archiv für mikroskopische Anatomie, 35, 68–87.

Heinrich, B. (1995). Neophilia and exploration in juvenile common ravens, Corvus corax. Animal Behaviour, 50(3), 695–704.

Hill, G. E., Doucet, S. M., & Buchholz, R. (2005). The effect of coccidial infection on iridescent plumage coloration in wild turkeys. Animal Behaviour, 69(2), 387–394.

Hill, G. E. & McGraw, K. J., Eds. (2006). Bird coloration, volume I. Cambridge, MA: Harvard University Press.

Hu, D., Clarke, J. A., Eliason, C. M., Qiu, R., Li, Q., Shawkey, M. D., Zhao, C., D’Alba, L., Jiang, J., & Xu, X. (2018). A bony-crested Jurassic dinosaur with evidence of iridescent plumage highlights complexity in early paravian evolution. Nature Communications, 9(217).

Hwang, V., Stephenson, A. B., Barkley, S., Brandt, S., Xiao, M., Aizenberg, J., & Manoharan, V. N. (2021). Designing angle-independent structural colors using Monte Carlo simulations of multiple scattering. Proceedings of the National Academy of Sciences, 118(4), e2015551118.

Hwang, V., Stephenson, A. B., Magkiriadou, S., Park, J.-G., & Manoharan, V. N. (2020). Effects of multiple scattering on angle-independent structural color in disordered colloidal materials. Physical Review E, 101(1), 012614.

Igic, B., D’Alba, L., & Shawkey, M. D. (2018). Fifty shades of white: how white feather brightness differs among species. The Science of Nature, 105(3), 18.

Iskandar, J.-P., Eliason, C. M., Astrop, T., Igic, B., Maia, R., & Shawkey, M. D. (2016). Morphological basis of glossy red plumage colours. Biological Journal of the Linnean Society, 119(2), 477–487.

Jacobs, I. F., Osvath, M., Osvath, H., Mioduszewska, B., von Bayern, A. M. P., & Kacelnik, A. (2014). Object caching in corvids: Incidence and significance. Behavioural Processes, 102, 25–32.

Jitsumori, M. & Delius, J. D. (2001). Object Recognition and Object Categorization in Animals. In T. Matsuzawa (Ed.), Primate Origins of Human Cognition and Behavior (pp. 269–293). Tokyo: Springer Japan.

Jakel, F., Singh, M., Wichmann, F. A., & Herzog, M. H. (2016). An overview of quantitative approaches in Gestalt perception. Vision Research, 126, 3–8.

Kinoshita, S., Yoshioka, S., & Miyazaki, J. (2008). Physics of structural colors. Reports on Progress in Physics, 71(7), 076401.

Komatsu, H., Nishio, A., Okazawa, G., & Goda, N. (2013). Computational Color Imaging. In S. Tominaga, R. Schettini, & A. Tŕemeau (Eds.), ‘Yellow’ or ‘Gold’?: Neural Processing of Gloss Information (pp. 1–12). Berlin, Heidelberg: Springer.

Loyau, A., Gomez, D., Moureau, B., Thery, M., Hart, N. S., Jalme, M. S., Bennett, A. T. D., & Sorci, G. (2007). Iridescent structurally based coloration of eyespots correlates with mating success in the peacock. Behavioral Ecology, 18(6), 1123–1131.

Magkiriadou, S., Park, J.-G., Kim, Y.-S., & Manoharan, V. N. (2014). Absence of red structural color in photonic glasses, bird feathers, and certain beetles. *Phys*. Rev. E, 90(6), 062302.

Maia, R., Caetano, J. V. O., Bao, S. N., & Macedo, R. H. (2009). Iridescent structural colour production in male blue-black grassquit feather barbules: the role of keratin and melanin. Journal of The Royal Society Interface, 6, S203–S211.

Maia, R., D’Alba, L., & Shawkey, M. D. (2011). What makes a feather shine? A nanostructural basis for glossy black colours in feathers. Proceedings of the Royal Society B: Biological Sciences, 278(1714), 1973–1980.

Maia, R., Eliason, C. M., Bitton, P. P., Doucet, S. M., & Shawkey, M. D. (2013). pavo: An R package for the analysis, visualization and organization of spectral data. Methods in Ecology and Evolution, 4(10), 906–913.

Matsumoto, T., Fukuda, K., & Uchikawa, K. (2015). Appearance of ‘gold’ affects glossiness and metallicity of a surface. Journal of Vision, 15(12), 819–819.

McMahon, B. C. (2017). Iridescence, vision, and belief in the Early Modern Hispanic World. PhD thesis, University of Southern California.

Meadows, M. G., Morehouse, N. I., Rutowski, R. L., Douglas, J. M., & McGraw, K. J. (2011). Quantifying iridescent coloration in animals: a method for improving repeatability. Behavioral Ecology and Sociobiology, 65(6), 1317–1327.

Meert, K., Pandelaere, M., & Patrick, V. M. (2014). Taking a shine to it: How the preference for glossy stems from an innate need for water. Journal of Consumer Psychology, 24(2), 195–206.

Michelson, A. (1911). On metallic colouring in birds and insects. *The London*, Edinburgh, and Dublin Philosophical Magazine and Journal of Science, 21(124), 554–567.

Montgomerie, R. (2006). Analyzing Colors. In K. J. Mcgraw & G. E. Hill (Eds.), Bird Coloration Vol. I: Mechanisms and Measurements (pp. 90–147). Cambridge, MA: Harvard University Press.

Noh, H., Liew, S. F., Saranathan, V., Mochrie, S. G. J., Prum, R. O., Dufresne, E. R., & Cao, H. (2010a). How Noniridescent Colors Are Generated by Quasi-ordered Structures of Bird Feathers. Advanced Materials, 22(26-27), 2871–2880.

Noh, H., Liew, S. F., Saranathan, V., Prum, R. O., Mochrie, S. G. J., Dufresne, E. R., & Cao, H. (2010b). Contribution of double scattering to structural coloration in quasiordered nanostructures of bird feathers. Physical Review E, 81(5), 051923.

Noh, H., Liew, S. F., Saranathan, V., Prum, R. O., Mochrie, S. G. J., Dufresne, E. R., & Cao, H. (2010c). Double scattering of light from Biophotonic Nanostructures with short-range order. Optics Express, 18(11), 11942–11948.

Norden, K. K., Eliason, C. M., & Stoddard, M. C. (2021). Evolution of brilliant iridescent feather nanostructures. eLife, 10, e71179.

Norden, K. K., Faber, J. W., Babarovic, F., Stubbs, T. L., Selly, T., Schiffbauer, J. D., Stefanic, P. P., Mayr, G., Smithwick, F. M., & Vinther, J. (2019). Melanosome diversity and convergence in the evolution of iridescent avian feathers—Implications for paleocolor reconstruction. Evolution, 73(1), 15–27.

Norman, J. F., Todd, J. T., & Phillips, F. (2020). Effects of illumination on the categorization of shiny materials. Journal of Vision, 20(5), 2.

Okazawa, G., Goda, N., & Komatsu, H. (2012). Selective responses to specular surfaces in the macaque visual cortex revealed by fMRI. NeuroImage, 63(3), 1321–1333.

Osorio, D. & Cuthill, I. C. (2015). Camouflage and perceptual organization in the animal kingdom. In J. Wagemans (Ed.), The Oxford handbook of perceptual organisation. Oxford University Press.

Osorio, D. & Ham, A. D. (2002). Spectral reflectance and directional properties of structural coloration in bird plumage. Journal of Experimental Biology, 205(14), 2017–2027.

Ospina-Rozo, L., Roberts, A., & Stuart-Fox, D. (2022). A generalized approach to characterize optical properties of natural objects. Biological Journal of the Linnean Society, 137(3), 534–555.

Ozaki, R., Kikumoto, K., Takagaki, M., Kadowaki, K., & Odawara, K. (2021). Structural colors of pearls. Scientific Reports, 11(1), 15224.

Parker, A. R., Mckenzie, D. R., & Large, M. C. J. (1998). Multilayer Reflectors in Animals Using Green and Gold Beetles as Contrasting Examples. Journal of Experimental Biology, 201(9), 1307–1313.

Prum, R. O. (2006). Anatomy, physics, and evolution of structural colors. In G. E. Hill & K. J. McGraw (Eds.), Bird coloration (pp. 295–353). Cambridge, MA: Harvard University Press.

Prum, R. O., Dufresne, E. R., Quinn, T., & Waters, K. (2009). Development of colour-producing *β*-keratin nanostructures in avian feather barbs. Journal of The Royal Society Interface, 6, S253–S265.

Quintero, E. & Espinosa de los Monteros, A. (2011). Microanatomy and evolution of the nanostructures responsible for iridescent coloration in Trogoniformes (Aves). Organisms Diversity & Evolution, 11(3), 237.

Rump, M., Muller, G., Sarlette, R., Koch, D., & Klein, R. (2008). Photo-realistic Rendering of Metallic Car Paint from Image-Based Measurements. Computer Graphics Forum, 27(2), 527–536.

Saranathan, V., Forster, J. D., Noh, H., Liew, S.-F., Mochrie, S. G. J., Cao, H., Dufresne, E. R., & Prum, R. O. (2012). Structure and optical function of amorphous photonic nanostructures from avian feather barbs: a comparative small angle X-ray scattering (SAXS) analysis of 230 bird species. Journal of The Royal Society Interface, 9(75), 2563–2580.

Schmid, A. C., Barla, P., & Doerschner, K. (2023). Material category of visual objects computed from specular image structure. Nature Human Behaviour, 7(7), 1152– 1169.

Schumacher, S., Burt de Perera, T., Thenert, J., & von der Emde, G. (2016). Cross-modal object recognition and dynamic weighting of sensory inputs in a fish. Proceedings of the National Academy of Sciences, 113(27), 7638–7643.

Seago, A. E., Brady, P., Vigneron, J.-P., & Schultz, T. D. (2009). Gold bugs and beyond: a review of iridescence and structural colour mechanisms in beetles (Coleoptera). Journal of The Royal Society Interface, 6, S165–S184.

Sekuler, R., Watamaniuk, S. N. J., & Blake, R. (2002). Perception of visual motion. In Steven’s handbook of experimental psychology: Sensation and perception, Vol. 1, 3rd ed (pp. 121–176). Hoboken, NJ, US: John Wiley & Sons Inc.

Shawkey, M. D., Hauber, M. E., Estep, L. K., & Hill, G. E. (2006). Evolutionary transitions and mechanisms of matte and iridescent plumage coloration in grackles and allies (Icteridae). Journal of The Royal Society Interface, 3(11), 777–786.

Shawkey, M. D. & Hill, G. E. (2005). Carotenoids need structural colours to shine. Biology Letters, 1(2), 121–124.

Shephard, T. V., Lea, S. E. G., & Hempel de Ibarra, N. (2015). ‘The thieving magpie’? No evidence for attraction to shiny objects. Animal Cognition, 18(1), 393–397.

Silvia, P. J., Christensen, A. P., Cotter, K. N., Jackson, T. A., Galyean, C. B., Mc-Croskey, T. J., & Rasheed, A. Z. (2018). Do people have a thing for bling? Examining aesthetic preferences for shiny objects. Empirical Studies of the Arts, 36, 101–113.

Silvia, P. J., Rodriguez, R. M., Cotter, K. N., & Christensen, A. P. (2021). Aesthetic Preference for Glossy Materials: An Attempted Replication and Extension. Behavioral Sciences, 11(4), 44.

Simpson, J. A., Weiner, E. S. C., & Press, O. U., Eds. (1989). The Oxford English dictionary. Oxford: Oxford; New York: Clarendon Press; Oxford University Press, 2nd ed edition.

Simpson, R. K. & McGraw, K. J. (2019). Interspecific Covariation in Courtship Displays, Iridescent Plumage, Solar Orientation, and Their Interactions in Hummingbirds. The American Naturalist, 194(4), 441–454.

Skigin, D. C., Inchaussandague, M. E., D’Ambrosio, C., Barreira, A., & Tubaro, P. (2019). How the observed color of the Swallow Tanager (Tersina viridis) changes with viewing geometry. Optik, 182, 639–646.

Soto, F. A. & Wasserman, E. A. (2014). Mechanisms of object recognition: what we have learned from pigeons. Frontiers in Neural Circuits, 8.

Stavenga, D. G., Leertouwer, H. L., Osorio, D. C., & Wilts, B. D. (2015). High refractive index of melanin in shiny occipital feathers of a bird of paradise. Light: Science and Applications, 4(1), e243.

Stavenga, D. G., Leertouwer, H. L., & Wilts, B. D. (2018). Magnificent magpie colours by feathers with layers of hollow melanosomes. Journal of Experimental Biology, 221(4).

Stavenga, D. G., van der Kooi, C. J., & Wilts, B. D. (2017). Structural coloured feathers of mallards act by simple multilayer photonics. Journal of The Royal Society Interface, 14(133), 20170407.

Stoddard, M. C. & Prum, R. O. (2008). Evolution of Avian Plumage Color in a Tetrahedral Color Space: A Phylogenetic Analysis of New World Buntings. The American Naturalist, 171(6), 755–776.

Stuart-Fox, D., Ospina-Rozo, L., Ng, L., & Franklin, A. M. (2020). The Paradox of Iridescent Signals. Trends in Ecology & Evolution.

Sumner, R. (2014). Processing RAW images in MATLAB.

Sutton, P. & Snow, M. (2015). Iridescence: The Play of Colours. Thames & Hudson Australia.

Thomas, D. B., McGraw, K. J., Butler, M. W., Carrano, M. T., Madden, O., & James, H. F. (2014). Ancient origins and multiple appearances of carotenoid-pigmented feathers in birds. Proceedings of the Royal Society B: Biological Sciences, 281(1788), 20140806–20140806.

Thomas, D. H. N., Kjernsmo, K., Scott-Samuel, N. E., Whitney, H. M., & Cuthill, I. C. (2023). Interactions between color and gloss in iridescent camouflage. Behavioral Ecology, (pp. arad050).

Todd, J. T. & Norman, J. F. (2018). The visual perception of metal. Journal of Vision, 18(3), 9–9.

Urquia, G. M., Inchaussandague, M. E., Skigin, D. C., Lester, M., Barreira, A., & Tubaro, P. (2020). Theoretical approaches to study the optical response of the red-legged honeycreeper’s plumage (Cyanerpes cyaneus). Applied Optics, 59(13), 3901–3909.

Wagemans, J., Elder, J. H., Kubovy, M., Palmer, S. E., Peterson, M. A., Singh, M., & von der Heydt, R. (2012). A century of Gestalt psychology in visual perception: I. Perceptual grouping and figure–ground organization. Psychological Bulletin, 138, 1172–1217.

Wilts, B. D., Michielsen, K., Raedt, H. D., & Stavenga, D. G. (2014). Sparkling feather reflections of a bird-of-paradise explained by finite-difference time-domain modeling. Proceedings of the National Academy of Sciences, 111(12), 4363–4368.

